# Network-based modelling reveals cell-type enriched patterns of non-coding RNA regulation during human skeletal muscle remodelling

**DOI:** 10.1101/2024.08.11.606848

**Authors:** Jonathan C. Mcleod, Changhyun Lim, Tanner Stokes, Jalil-Ahmad Sharif, Vagif Zeynalli, Lucas Wiens, Alysha C D’Souza, Lauren Colenso-Semple, James McKendry, Robert W. Morton, Cameron J. Mitchell, Sara Y. Oikawa, Claes Wahlestedt, J Paul Chapple, Chris McGlory, James A. Timmons, Stuart M. Phillips

**Author notes:** Correspondence to: Jonathan C. Mcleod. Joint senior authors (James A. Timmons, Stuart M. Phillips).

## Abstract

A majority of human genes produce non-protein-coding RNA (ncRNA), and some have roles in development and disease. Neither ncRNA nor human skeletal muscle is ideally studied using short-read sequencing, so we used a customised RNA pipeline and network modelling to study cell-type specific ncRNA responses during muscle growth at scale. We completed five human resistance-training studies (n=144 subjects), identifying 61% who successfully accrued muscle-mass. We produced 288 transcriptome-wide profiles and found 110 ncRNAs linked to muscle growth *in vivo,* while a transcriptome-driven network model demonstrated interactions via a number of discrete functional pathways and single-cell types. This analysis included established hypertrophy-related ncRNAs, including *CYTOR* – which was leukocyte-associated (FDR = 4.9 x10^-7^). Novel hypertrophy-linked ncRNAs included *PPP1CB-DT* (myofibril assembly genes, FDR = 8.15 x 10^-8^), and *EEF1A1P24* and *TMSB4XP8* (vascular remodelling and angiogenesis genes, FDR = 2.77 x 10^-5^). We also discovered that hypertrophy lncRNA *MYREM* shows a specific myonuclear expression pattern *in vivo*. Our multi-layered analyses established that single-cell-associated ncRNA are identifiable from bulk muscle transcriptomic data and that hypertrophy-linked ncRNA genes mediate their association with muscle growth via multiple cell types and a set of interacting pathways.

**One Sentence Summary:** We used an optimised transcriptomic strategy to identify a set of ncRNA genes regulated during skeletal muscle hypertrophy in one hundred and forty-four people, with network modelling and spatial imaging providing biological context.

## INTRODUCTION

Most genes in the human genome do not code for a protein ^1^ but instead are transcribed into non-coding RNAs (ncRNA), including long non-coding RNAs (lncRNA)^2^. Early work demonstrated that the proportion of ncRNA genes increases with organismal complexity ^3^, and it is now recognised that ncRNAs participate in many key biological roles ^4,5^, including regulating genome organisation and gene expression through nucleic and RNA–protein interactions ^4,6^. Further, ncRNAs can act as a surveillance mechanism to prevent the accumulation of aberrant products of protein translation ^7^, participate in epigenetic regulation ^8,9^, and several ncRNAs are directly implicated in human diseases ^6,10–13^. While ncRNAs classically have no-protein coding potential, emerging evidence suggests ncRNAs can harbour micro-peptides that have biological activity ^14^. Nevertheless, most ncRNAs have no characterised function and are not reliably detected in human muscle using short-read sequencing ^15^, highlighting a need for their accurate measurement and biological characterisation.

Skeletal muscle accounts for ∼40% of total body mass ^16^, is paramount in various mechanical and metabolic functions and is subject to age-related decline ^17^. Resistance exercise training (RT) results in muscle fibre hypertrophy, mitigating ageing-related loss of muscle ^18^. Yet not all humans robustly accrue skeletal muscle mass, even following prolonged supervised RT ^19^. This lack of response is indicative that the influence of RT on growth is often not readily discernable beyond the technical variation of a measurement instrument ^20^. Although exogenous factors – such as RT load or frequency ^21^ and ensuring adequate dietary protein provision ^22^ – modestly contribute to the load-induced hypertrophic phenotype, the intrinsic genomic characteristics of an individual are hypothesised to determine the bulk of the variability in hypertrophic response ^23^. Early human studies stratifying RT-induced hypertrophy identified a paradoxical gene signature consistent with less mammalian target of rapamycin complex 1 (mTORC1) activation in individuals with the greatest hypertrophy ^24^ or altered microRNA response profiles ^19^. Ultimately, studying individuals with the most robust responses to muscle loading may yield novel insights into causal regulators of adaptation and potentially identify therapeutic targets for treating muscle loss, including sarcopenia ^25^.

The current understanding of exercise transcriptomics is mostly limited to protein-coding genes; however, ncRNAs have begun to emerge as critical regulators of skeletal muscle growth. Recently, Wohlwend and colleagues ^26^ identified the lncRNA cytoskeleton regulator RNA (*CYTOR*) as regulated during muscle growth, negatively regulated with age and one that promoted fast skeletal muscle fibre expression in mice by sequestering the TEA domain transcription factor 1 (*TEAD1*). We discovered that the lncRNA, *FRAIL1*, was increased with muscle age ^27^, and more recently, increased *FRAIL1* was found to negatively influence muscle strength and function ^28^. Additionally, *Chronos* is a skeletal muscle-enriched lncRNA that was shown to repress hypertrophic growth in vitro and in vivo by negatively regulating Bmp7 expression ^29^. Despite the emerging recognition that ncRNAs act as regulators of skeletal muscle development ^30^, myogenesis, and musculoskeletal disease ^31^, most ncRNAs have no known function in adult muscle ^1^. The multi-nucleated and fibrous nature of skeletal muscle fibres makes it challenging to study using single-cell transcriptomics, and most single-cell technologies do not readily capture the noncoding transcriptome. In the present study, we generated five independent supervised exercise training studies (n=144 subjects) and applied our customised bulk RNA profiling strategy ^13,27,32^, which more reliably detects ncRNAs in human muscle ^15^. Combined with pathway analyses ^33,34^, data-driven quantitative RNA network modelling^35^ and selective spatial profiling, we were able to localise ncRNAs to specific cell types within muscle, providing insight into their biological role ^36,37^.

## METHODS

### Description of five independent clinical studies

For a brief overview of the analysis, please refer to Figure 1. Each exercise intervention study utilised *supervised* training to load the thigh muscle and promote hypertrophy. It is known that a wide range of exercise protocols yield similar group average increases in muscle mass ^20,21,24,38,39^. Accordingly, despite variations in training experience and training load, the relative gains in lean mass in each study demonstrated the same average increase. Note that the global transcriptome profiles were partially utilised (other than 50 new transcriptome profiles) to validate a list of 141 protein-coding genes identified during the remodelling of lean mass with altered loading ^25^. All five studies were approved by local research ethics boards and complied with all ethical standards for research involving human participants set by the Declaration of Helsinki.

**Figure 1.**
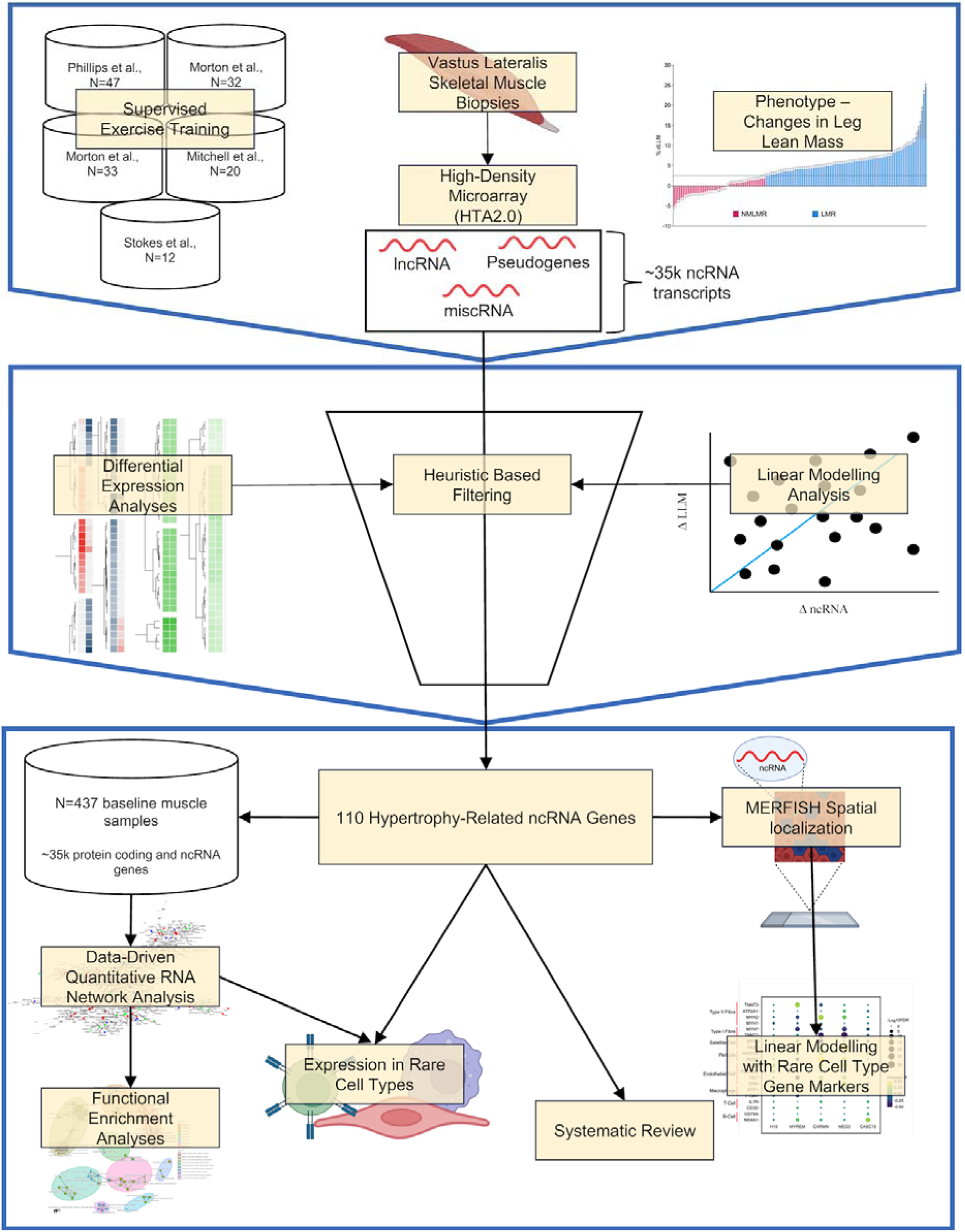
Overview of the analysis pipeline. We used 144 participants from five supervised muscle-loading studies. We measured skeletal muscle hypertrophy (using DXA and MRI), and 88 individuals exhibited a change in skeletal muscle hypertrophy beyond the technical variation of our measurement instruments. RNA was extracted from skeletal muscle samples pre- and post-training, and we quantified ncRNA transcriptome (miscRNA, lncRNA and pseudogenes) using the Human Transcriptome Array 2.0. Using differential expression analyses, linear modelling analysis, and heuristic-based filtering, we established 110 ncRNA genes are related to skeletal muscle hypertrophy. We used RNA network modelling analysis in baseline bulk skeletal muscle samples (n=437) of protein-coding and ncRNA genes. We used a novel transcript spatial localisation method to help facilitate the discovery of functional pathways and cell-type specific associations that several key ncRNA genes belong to.

### Study 1

Morton et al. ^40^ used recreationally active men (age: 22 ± 3 years; body mass index [BMI]: 26 ± 7 kg/m^2^). Each participant’s dominant leg was randomly assigned to a high-load, low-repetition (8-12 repetitions at ∼70-80% 1-repetition maximum [1RM]), RT protocol or a low-load, high-repetition (20-25 repetitions at ∼30-40% 1RM) RT protocol. The contralateral leg was assigned to the opposite condition providing two distinct interventions. Participants underwent RT three days a week (Monday, Wednesday, and Friday) for 10 weeks. Each RT session consisted of 3 sets of unilateral knee extensions to volitional fatigue (the starting leg alternated between RT sessions). RT-induced skeletal muscle hypertrophy did not differ between allocation conditions. Participants also received 25 g of whey protein isolate supplement twice daily (morning or post-exercise and pre-sleep) for the duration of the study. A muscle biopsy was obtained at pre- and post-training for 32 participants. Unilateral leg lean mass (LLM) was quantified using dual-energy X-ray absorptiometry (DXA) pre- and post-training.

### Study 2

Muscle biopsies originated from Morton et al. ^39^. Participants were active, with 4 – 5 years of RT experience, and then performed RT 4 days/week (Mon, Tues, Thurs, Fri) for 12 weeks (all the major muscle groups). Each session included 5 exercises performed for 3 sets to volitional failure. Participants were randomly assigned to complete either low-load, high-repetition (20-25 repetitions per set at ∼ 30% - 50% 1-RM) RT or high-load, low-repetition RT (8-12 repetitions per set at ∼ 75%-90% 1-RM). Indices of muscle hypertrophy did not differ between the training groups. Participants consumed 30 g of protein after each exercise session. Biopsy tissue was available at baseline and following 12 weeks of RT for 33 participants. The average of bilateral LLM was assessed using DXA at pre and post training.

### Study 3

Muscle biopsies originated from Phillips et al. ^41^. Sedentary participants performed high-intensity cycle training 3 days per week (Monday, Wednesday, and Friday) for 6 weeks. Each session consisted of a 2 min warmup at 50 W followed by 5 sets of high-intensity cycling at 125% VO_2 peak_ for 1 minute. Each set was separated by 90 s of rest. Biopsy tissue was available at pre- and post-training for 47 participants, including 18 males and 29 females with an average age of 39 years (range 21-51 years) and a BMI of 31 kg/m^2^ (range 26 - 43 kg/m^2^). Unilateral upper LLM was assessed using DXA at baseline and 3 days following the final training session.

### Study 4

Muscle biopsies originated from Mitchell et al. ^42^. Young, recreationally active men participated in whole-body RT 4 days per week for 16 weeks. RT sessions consisted of two upper-body and two lower-body training sessions per week. The program progressed from 3 sets of 12 repetitions to 4 sets of 6 repetitions of each exercise. The last set of each exercise was performed to volitional failure. Participants consumed 30 g of protein immediately after each exercise session. Biopsy tissue was available for 20 participants pre- and post-training. Unilateral thigh muscle volume was measured using Magnetic Resonance Imaging (MRI) pre- and post-training.

### Study 5

Muscle biopsies originated from Stokes et al. ^25^. Recreationally active men (age: 21 ± 3 years; BMI: 24 ± 3 kg/m^2^) underwent unilateral leg extension and leg press RT thrice weekly (Monday, Wednesday, and Friday) for 10 weeks. Specifically, each session consisted of 3 sets of 8-12 repetitions of leg extension and 3 sets of 8-12 repetitions on a leg press. The last set of RT was performed to volitional failure. Following each exercise session, participants ingested 25 g of whey protein isolate. Biopsy tissue was available for 12 participants pre- and post-training. Unilateral LLM was measured pre- and post-training using DXA.

### Defining hypertrophy beyond typical laboratory variation

Changes in DXA LLM (or, in one case, quadriceps muscle volume by MRI) defined subjects that demonstrated a genuine increase in muscle mass (‘responders’). For the remaining subjects, we could not show they demonstrated training-induced hypertrophy (we do not claim they have no response). We used the technical variation threshold for DXA-derived changes in LLM (2.0 – 2.4%)^43,44^ and MRI-derived changes in thigh muscle volume (2.3%)^45^ as cut-off values for these two groups. For all individuals, the percentage change in LLM (or quadriceps muscle volume for MRI) post-exercise training was calculated. Critically, there was no association between baseline LLM and changes in LLM (Figure S1; Table S1). Individuals who exhibited a ≥ 2.5% increase in LLM were defined as “LM Responders (LMR),” whereas individuals who showed < 2.0% change in LLM (or change in quadriceps muscle volume for ^42^) were defined as having “No measurable LM response (NMLMR).” These two groups (LMR N=88; NMLMR N=50) were studied separately for evidence of differential noncoding gene expression (DE). Only 6 individuals fell within the range of reported precision error (2.0 – 2.4%) of the DXA (or MRI), and they were included in the linear modelling analysis.

### Clinical data for quantitative network modelling

We used n=437 transcriptome profiles from mainly sedentary subjects, aged 50y or younger, to generate a human muscle tissue gene co-expression model using multiscale embedded gene co-expression network analysis (MEGENA)^35^. A total of 35,079 protein-coding and non-coding genes were used as input, and the analysis time was *∼600 hours* on a 3.2GHz 16-core Intel Xeon W-based Mac Pro (768GB 2933 MHz DDR4 RAM). Unlike most popular and fast network methods, MEGENA applies statistical thresholds to identify significant relationships at both the gene and network structure level, e.g. false discovery rate (FDR) < 1% for gene-gene Pearson correlation coefficients, P < 0.05 for module significance, P < 0.01 for network connectivity, with 1000 permutations for adjusted values (FDR). We present the overall hierarchical organisation of the planar filtered networks using a sunburst plot, and individual module plots were produced using Fruchterman-Reingold force-directed plotting within MEGENA ^35^. Gene co-expression modules that contained our candidate ncRNA genes were labelled in the sunburst plot and studied further (See below).

### RNA extraction and transcriptome profiling

In each case, RNA was extracted from approximately 20 mg of muscle tissue^25,39,40,42^ in 1000 mL of TRIzol, which was added to Lysing Matrix D tubes containing ceramic microbeads (MP Biomedicals, Solon, OH, USA) and homogenised using a FastPrep tissue homogeniser (MP Biomedicals, Solon, OH, USA). 200 mL of chloroform was added, and the tubes were hand-shaken vigorously for 15 s and incubated at room temperature for 5 min. Samples were then centrifuged at 12 000 g for 10 min at 4°C, and the upper aqueous phase containing RNA was transferred to an RNase-free tube. RNA was purified using the EZNA Total RNA Isolation kit (Omega Bio-Tek, Norcross, GA, USA). RNA was processed for transcriptome profiling using the GeneChip WT Plus Reagent Kit (no PCR) by processing to single-stranded sense DNA via a *reverse transcription linear priming strategy* that primes the poly-A and non-poly-A RNA. First and second-strand cDNA synthesis was performed using ∼100 ng of RNA and a spike-in Poly-A control, followed by reverse transcription into cRNA. cRNA was purified using magnetic beads and quantified using spectrophotometry (Nanodrop UV-Vis, Thermo Fisher Scientific). 15 mg of cRNA was then amplified and hydrolysed using RNase H (leaving single-stranded cDNA) and purified with magnetic beads. cDNA (5.5 mg) was then fragmented, labelled, and hybridised to an array. The array was washed and stained using a GeneChip Fluidics Station 450 (Thermo Fisher) and scanned using a GeneChip Scanner 3000 7G (McMaster Genomic Core Facility, Hamilton). Samples from Phillips et al. ^41^ were processed using a Qiagen Tissuelyser and processed as above without the additional purification step (Jensen Laboratory, MPI, Horsholm, Denmark).

### Computational processing of the HTA 2.0 array platform

We updated our 2017 process for annotating and processing the transcriptomic technology^13,15,25,27,46^. Briefly, the HTA 2.0 array has 6.9 million 25-mer probes and the annotation of each probe is checked against the recent reference genome and transcriptomes^47,48^ using R. To do this, we produce a standardised FASTA (with a unique label for each 25mer probe) file, a probe GC content file and a probe-level chip definition file (CDF). For the present study, the FASTA file was aligned against Grch38 - Gencode 43 ^49^ using the STAR aligner ^50^. Probes that uniquely map were combined to form groups of probes (probe-sets), and each probe-set relied on n>3 probes (median = 66). In addition, the raw probe signal of a large number of diverse human muscle profiles (n=1124, GSE154846) was analysed using the probe-level CDF and aroma.affymetrix ^47^, affy ^51^ and the affxparser (bioconductor.org/packages/affxparser/) packages. Probes with both a very low signal and a low coefficient of variation were removed to arrive at a final set of probes to be included in the CDF^13,15,25,27,46^. This CDF was then used to summarise RNA expression from the GC-corrected CEL files produced from Studies 1-5 and 437 pre-existing CEL files, after which iterative rank-order normalisation (IRON) ^52^ in the default mode was used to generate a normalised expression matrix. From the starting 6.9M probes, reannotation and summarisation yielded a transcript level probe-set CDF used 5.23M probes to define 220.6K probe-sets prior to study-specific transcript signal filtering (Supplement Data S1). The custom CDF used in this study is deposited at GSE154846, along with new raw data produced for this study (GSE270823). This updated CDF can be utilised to extract any other human muscle data sets produced on the HTA 2.0 platform. We estimated that a total of 173,594 transcripts were annotated above the background signal using standard deviation-based filtering.

### Differential expression analyses of ncRNA genes

For DE analyses, we used SAMR ^53^ and 10K permutations to estimate the false discovery rate (FDR), a method that appears more robust than other p-value correction methods ^54^. Paired SAMR analyses were applied to the LMR group (176 paired RNA samples) and the NLMR group (100 paired RNA samples). We relied on an FDR cut-off of 5% and a ≥ |1.2| fold change to generate differentially expressed ncRNA lists. To focus on the most distinct observations between the LMR and NLMR groups, we applied the following heuristics. We prioritised ncRNA transcripts that displayed a fold change in opposite directions across our two response groups or where the differentially expressed ncRNA showed a fold change ratio between the groups ≥ |1.2|. Finally, to clarify if there were any ncRNAs differentially expressed at baseline between our groups (note baseline physiology could not distinguish LMR and NLMR groups), we utilised an unpaired SAMR analysis (using the same criteria for a differentially expressed transcript as described above).

### Assessing the relationship between changes in gene expression and changes in lean mass

The linear relationship between changes in ncRNA transcript expression and changes in LLM (or change in quadriceps muscle volume for ^42^) was examined using a type III analysis of variance (R Core Team (2021). R: A language and environment for statistical computing. URL https://www.R-project.org/.). We used the following linear model: changes in ncRNA transcript ∼ changes in LLM. A Pearson correlation coefficient (r) and a p-value were calculated for each transcript per study. Using the three largest cohort studies ^39–41^, we combined p-values using the Stouffer method, using the poolr R package ^55^. For transcripts that passed filters 1 and 2 (see below), FDRs were calculated ^56^ using multtest ^57^. ncRNA transcripts that were deemed to change linearly in proportion with delta lean mass passed all of the following filters: 1) 5-study median r (absolute) ≥0.2; 2) consistent (direction) r value in 4/5 studies; and 3) an FDR < 10% (the median was FDR 8%).

### Cell type-specific gene markers and MERSCOPE-based transcript localisation

We calculated Pearson correlation coefficients and p-values (FDR adjusted) to establish the association of ncRNAs or gene co-expression modules with markers of type I and type II muscle fibres and rarer mononuclear cell types found in bulk skeletal muscle (satellite cells, endothelial cells, pericytes, macrophages, T-cells, and B-cells). These analyses relied on a network model generated using 437 muscle biopsy samples in individuals aged 50y or younger (GSE154846). We used the following myonuclei and mononuclear cell-specific gene markers: type I fibres (*MYH7, TNNT1*)^58–60^, type II fibres (*MYH1, MYH2, ATP2A1, TNNT3*) ^58–60^, satellite cells (*PAX7, MYF5*) ^58,61^, endothelial cells (*ENG, TIE1, PECAM1, APLNR*), pericytes (*RGS5, HIGD1B*) ^58,62^, macrophages (*F13A1, SPP1*) ^58^, T cells (*CD3D, IL7R*) ^58,63^, and B cells (*MS4A1, CD79A*) ^58,63^. The MERSCOPE is a spatially resolved single-cell transcriptome profiling technology. As part of the process of designing and validating ongoing work in our laboratory, we produced images of the location of a small number of the ncRNAs identified in the present study. The MERSCOPE assay also included markers for muscle fibre types, satellite cells, and endothelial cells (type I fibre [MYH7 fish probe] ^58–60^, type II fibres [ATP2A1 fish probe] ^58–60^, satellite cells [PAX7, MYF5 merfish probes] ^58,61^ and endothelial cells [ENG, TIE1, PECAM1, APLNR merfish probes]). The panel was used to profile three independent muscle samples originating from the present study, and some ncRNA data pertinent to the present results is presented. The gene list was sent to Vizgen Inc, and they selected 30-50 merfish probes per gene for profiling on the MERSCOPE. Briefly, each fresh frozen sample embedded in the optimal cutting temperature compound (OCT, Tissue-Tek, The Netherlands) was sectioned at a thickness of 10 um in a cryostat at −20 °C and transferred onto the circular MERSCOPE glass slide (PN 2040001, Vizgen, USA) within the fiducial bead border, and prepared for transport ^64,65^. Tissue sections were fixed in pre-warmed 4% paraformaldehyde for 1 hour at 37°C. After fixation, the tissue sections were washed in 1X PBS 3 times for 5 min at room temperature, permeabilised in 70% ethanol (molecular grade) at 4°C overnight. Then, Vizgen cell boundary staining kit (PN 10400009, Vizgen, USA) was applied to stain cell membranes on the tissue sections after a 1-hour incubation in Blocking Solution mixed with RNase inhibitor (PN 20300100, Vizgen, USA). Afterwards, the tissue sections were incubated with Formamide Wash Buffer (PN 20300002, Vizgen, USA) at 37°C for 30 min and incubated in the custom-designed merfish panel at 37°C for about 40h. Then, the slide was incubated in the Formamide Wash Buffer at 47°C (30 min ×2), rinsed with the sample preparation buffer, and stained with the DAPI and poly(T) Staining reagents, and the fiducial fluorescent beads (perimeter of scannable area). The probes were imaged using the MERSCOPE (Vizgen Core Laboratory, Boston, USA) with default settings (poly(T) and DAPI and Cell Boundary Channels “on”, while scan thickness was 10μm). The raw images were converted to “vzg” meta-output files. For ncRNAs included in both assays, we used the MERSCOPE visualiser software and cell-specific marker probes to evaluate the location of each ncRNA.

### Functional enrichment pathway analysis

Enrichment analyses (e.g., gene ontology) ^33^ are used to uncover the functional biology of lists of genes, categorising genes by their known biochemical functions ^66^. For each network module, enrichment analysis was executed using the web tool Database for Annotation, Visualisation, and Integrated Discovery (DAVID; https://david.ncifcrf.gov/) ^33^, versus a custom background file containing the protein-coding genes estimated to be expressed in skeletal muscle tissue in the present study ^66^. Most ncRNAs are not ontologically characterised and thus do not contribute to the enrichment analyses. Data-driven RNA network modelling methods are constructed based on experimentally derived gene co-expression similarities^35^, and so provide a solution to categorising ncRNAs using “guilt by association”. Gene co-expression modules with less than 500 genes and gene ontology (GO) terms with an FDR ≤ 5.0×10^-5^ were selected for further study. The gene lists from the ncRNA-containing modules, along with the background list of expressed genes, were analysed using the compareCluster function^67,68^ in the ClusterProfiler R package utlising the GOTERM_BP_ALL from DAVID. ClusterProfiler was used to carry out functional enrichment using over-representation analyses and combined with enrichment mapping^67^ to establish links between distinct ncRNA-containing MEGENA modules. Note that not all gene identifiers from each module are mapped to the databases being used to generate these pathway analyses (a common problem).

## RESULTS

### Skeletal muscle hypertrophy following supervised exercise training

The average change in LLM (using DXA) across ^25,39–41^ was 3.4 ± 3.9% (range: −4.9 % to 18.8%); the average change in quadriceps muscle volume using MRI ^42^ was 7.1 ± 7.0% (range: −1.8% to 24.7%). Out of 144 participants, 88 individuals (61% of the total sample) were defined as LMR, whereas 50 individuals (34% of the total sample) did not demonstrate LLM changes beyond typical measurement error (NMLMR group). Thus, the average change in LLM for the LMR group and the NMLMR group was 6.6 ± 3.9% (range: 2.5% to 24.7%) and −0.6 ± 1.8 % (range: −4.9% to 1.9%), respectively (Figure 2). Baseline LLM status could not explain the observed variation in LLM gain (Figure S1; the demographics of the individual studies can be found in Table S1). We do not claim that the NMLMR subjects can not grow muscle tissue, just that we are unable to unequivocally assign any change in LLM beyond previously established method variation and we treated them as phenotypically divergent on this basis.

**Figure 2.**
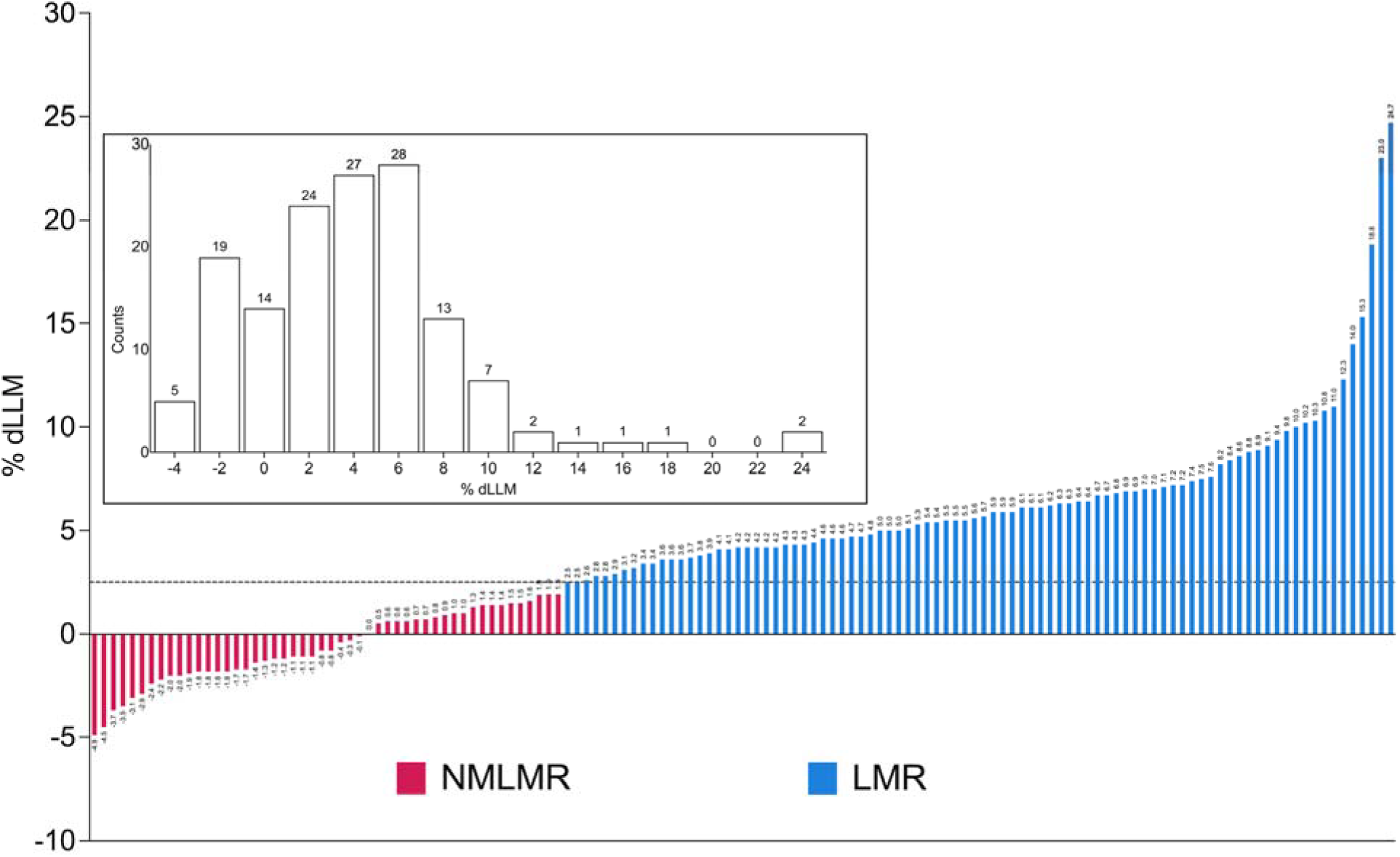
Phenotypic overview of included studies. Waterfall plot showing the percent change in leg lean mass (% dLLM) for 138 participants used for differential expression analyses. The dotted line (2.5% dLLM) depicts the threshold for the precision error of the DXA and MRI. Individuals who exhibited a ≥ 2.5% dLLM (or quadriceps muscle volume for MRI) were allocated to the group: “Lean Mass Responders” (LMR; n=88), whereas individuals who showed < 2.0% dLLM were assigned to the group called “No Measurable Lean Mass Response” (NMLMR; n=50). Each bar corresponds to one individual, and the number above each bar corresponds to the numerical value for dLLM%. Box Inset: Histogram depicts the number of individuals (144) falling into each %dLLM bin, and counts above each bar correspond to the number of individuals in that particular %dLLM bin. For a detailed overview of participant characteristics, please refer to Table S1.

### Overview of RNA molecules measured in bulk human skeletal muscle using high-density arrays

58% of quantified genes were protein-coding after filtering out low signal (n=13249, Figure 3A). The majority of detectable ncRNA genes in skeletal muscle are annotated as “lncRNA” molecules (7173 genes), which are broadly categorised as genes that are ≥ 200bp in length. Other classes of ncRNAs included miscellaneous RNA (n=1192; ncRNA genes that are <200bp in length) and pseudogenes (n=1254). Note that ncRNA genes are routinely described as ‘low-abundance’ genes ^36^ based on sequencing cDNA libraries. We find here that the expression profile of ncRNA genes is similar to protein-coding genes in human skeletal muscle (Figure 3B) using a library preparation method that does not rely on PCR and is not 3’ biased - consistent with a previous meta-analysis ^15^ from our group. The superior coverage with our array protocol over typical sequencing protocols most likely reflects, firstly, a more diverse cRNA library and, secondly, the lack of use of a competitive/counting quantification method, enabling more equitable detection of all RNA species (where array probes exist for that RNA – which in practice results in substantial greater coverage of the muscle transcriptome^15^).

**Figure 3.**
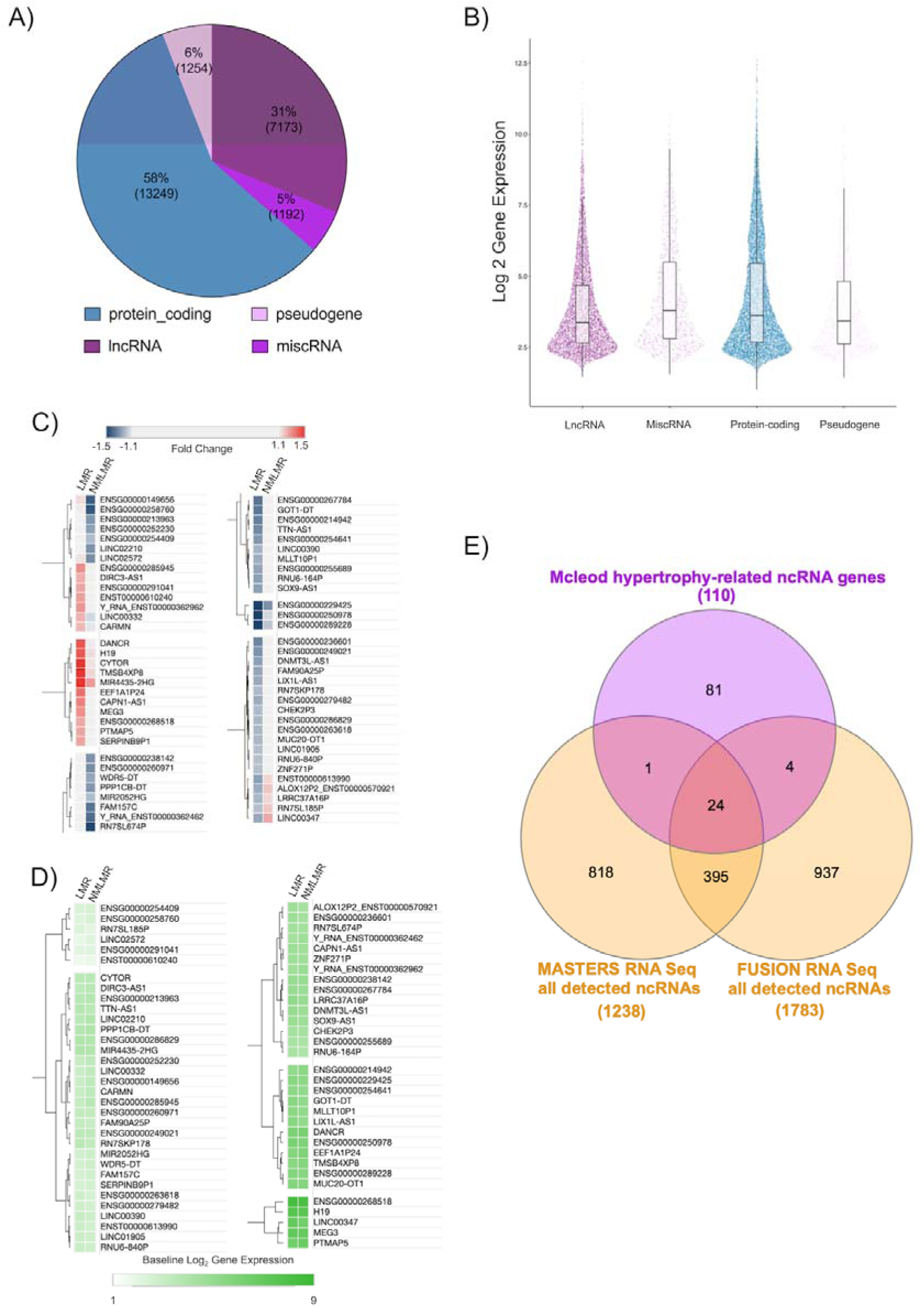
RNA molecules quantified on high-density arrays and establishing a 110 ncRNA gene signature associated with skeletal muscle hypertrophy. A) Quantitative overview of the number of genes (2 SD filter applied to protein-coding genes and 1 SD filter to ncRNA genes) belonging to each class of RNA. B) The distribution of the abundance (log 2 signal) of each RNA class is visualised using the median (horizontal line within the box), upper and lower interquartile ranges (upper and lower edges of the box), upper and lower whiskers (1.5 x interquartile range), and the points correspond to an individual gene. C) Heat plot that denotes fold changes for 65 ncRNA genes were differentially expressed and uniquely regulated in a manner that was distinct across the LMR group (50 genes) and the NMLMR group (15 genes). (D) The heat plot denotes the log2 baseline gene expression for the 65 uniquely regulated ncRNA genes. The heat plots (C, D) were created using Morpheus (https://clue.io/Morpheus), and genes were hierarchically clustered using Euclidean distance (linkage method: complete). E) 110 hypertrophy-related ncRNA genes were contrasted to all consistently detected ncRNA genes from two short-read RNA sequencing data sets profiling skeletal muscle (MASTERS ≈34 million reads, n = 136 ^72^; FUSION ≈60 million reads, n =278 ^71^ using a Venn diagram tool (https://www.interactivenn.net/; ^129^).

### Establishing a set of ncRNA genes related to skeletal muscle hypertrophy

One hundred ninety-three ncRNA genes were significantly regulated in the LMR group (Supplement Data, S2), whereas 50 genes were significantly regulated in the NMLMR group (Supplement Data, S3). To enable us to focus on the most robust results, we applied heuristic filtering (see Methods) to identify those ncRNA that demonstrated the most distinct pattern of regulation between our two groups^13,69,70^. This analysis revealed a subset of 50 ncRNA genes that were specifically regulated just in the LMR group and so may be directly linked with hypertrophy. Fifteen ncRNA genes were uniquely regulated in the NMLMR group (Figure 3C), and these may be associated with processes that impair growth or aim (unsuccessfully) to overcome negative influences on growth.

Notably, only two ncRNA genes were differentially expressed between the two groups prior to supervised training (Supplement Data S4), demonstrating that these baseline ncRNA transcriptomic profiles, much like the physiological characteristics, did not distinguish LMR from NMLMR groups (Figure 3D). In addition to DE analyses applied to the two post-hoc categorical groups (i.e., LMR group vs NMLMR group), linear analysis was used to determine if there was a linear relationship between changes in ncRNA gene expression and changes in LLM, using all 144 subjects. This analysis identified 46 ncRNA genes with a modest relationship to gain in LLM (median r= 0.26; median FDR = 8%; Figure S2A; Supplement Data S5). Combined with the DE analyses, a final list of 110 hypertrophy-related ncRNA genes (Supplement Data S6) was considered further. Notably, we identified most of the known muscle hypertrophy-related ncRNA genes (e.g. *CYTOR, MYREM, H19,* and *MEG3*), which now have some established biochemical roles in skeletal muscle (Table S2). The rest of the hypertrophy ncRNAs had no prior associated function in skeletal muscle (Table S2). Cardiac mesoderm enhancer-associated non-coding RNA (*CARMN)* was the only ncRNA gene that displayed a linear association with changes in LLM (Figure S2B-F) *and* was differentially expressed in the LMR group. Critically, only 30% of the hypertrophy ncRNA genes are consistently detected in skeletal muscle by RNA sequencing ^71,72^ (Figure 3E; Supplement Data S7), and only five ncRNA genes (*CASC15, MYREM, DANCR, MIR4435-2HG, MEG3*) were reported as being regulated by RT when using RNAseq ^72^.

### Spatial distribution of ncRNA genes in human skeletal muscle

There are several multi-plex true single-cell spatial technologies^64,65,73^, and we present here data from the MERSCOPE single-cell spatial workflow to provide the visual location of a few hypertrophy ncRNA genes, along with markers of type I and type II muscle fibres, satellite cells and endothelial cells ^58^. The prototype MERSCOPE assay (for an overview of the 3 samples used for MERSCOPE-MERFISH, please refer to Figure S3) showed reasonable agreement with the bulk expression data from the same biopsy sample (r = 0.78 – 0.81). In our bulk transcriptomic data, *MYREM* (*MYBPC2* cis regulating lncRNA enhancer of myogenesis (ENSG00000268518)) expression was positively associated with type II fibre markers (e.g., *TNNT3*: r =0.37; FDR = 2.57 x10^-14^; Figure 4A) ^58^, and negatively associated with type I fibre markers (e.g., *MYH7:* r = −0.33; FDR = 1.45 x10^-11^; Figure 4A) ^58^. Visually, *MYREM* is concentrated beneath the basal lamina and was not observed in nuclei with gene markers of satellite cells (*PAX7, MYF5*) or endothelial cells (*APLNR, TIE1, KDR, ENG*), indicating that *MYREM* is a novel marker of mature myofibril nuclei in humans (Figure 4B), especially in type II fibres. This interpretation is supported by recent data from Chang and colleagues ^74^ demonstrating that *MYREM* increases the transcriptional expression of *MYBPC2*, a type II isoform of skeletal muscle myosin-binding protein C ^75,76^.

**Figure 4.**
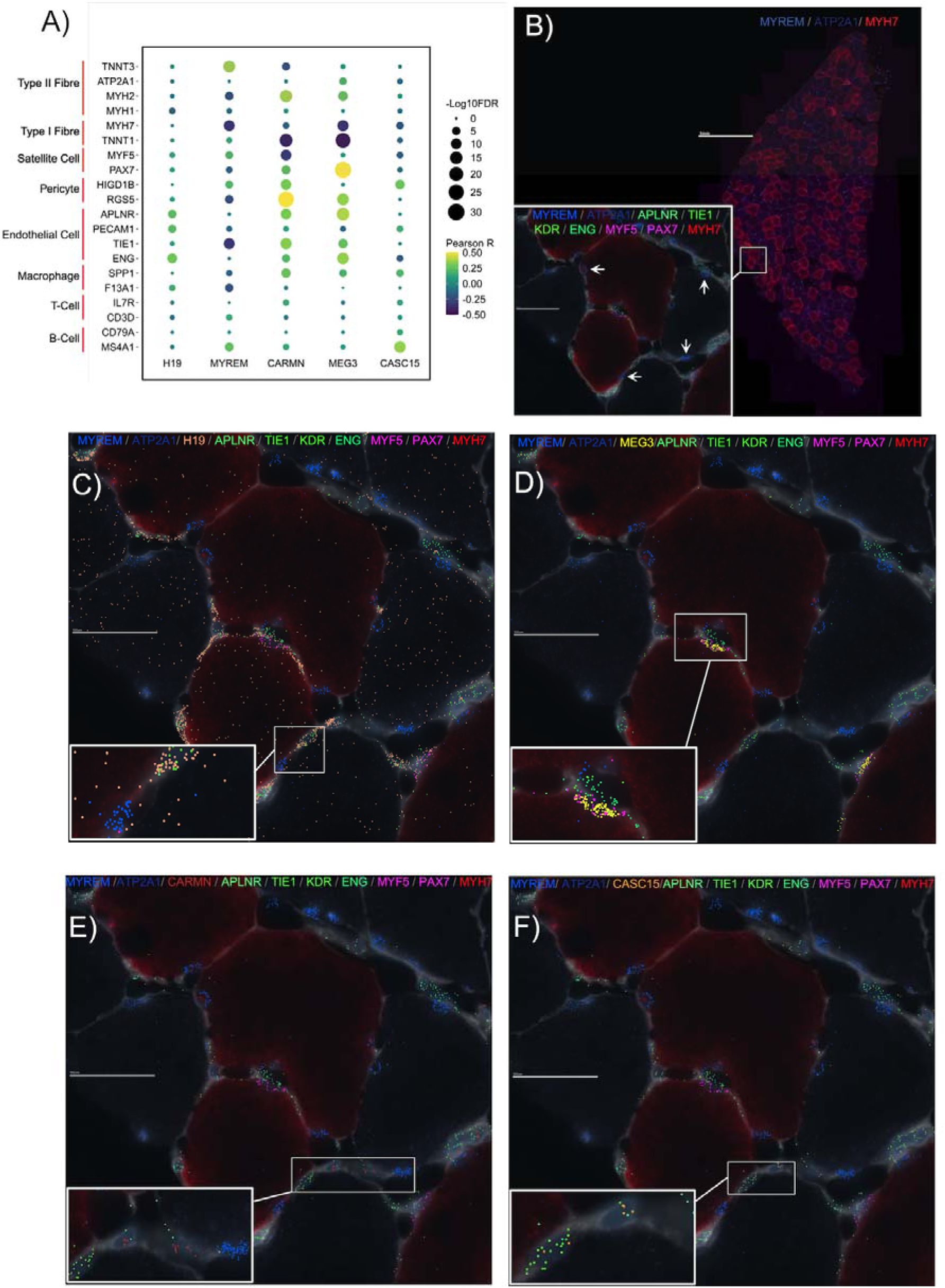
Spatially resolving several hypertrophy-related ncRNA genes in human skeletal muscle *in vivo*. A) Dot plot depicting the association between the expression of hypertrophy-related ncRNA genes - that were used for MERFISH - and gene markers for skeletal muscle fibre types and rare mononuclear cell types, using bulk transcriptomic data (n=437). The colour of the dot corresponds to the Pearson correlation coefficient, and the size of the dot is proportional to the –Log10 FDR. B) High-level image of type I (*MYH7*) and type II (*ATP2A1*) FISH probes on a human skeletal muscle cross-section. Box inset shows magnified region of skeletal muscle cross-section with MERFISH probes for *MYREM,* as well as MERFISH probes for endothelial cell protein-coding gene markers (*APLNR, TIE1, KDR, ENG*) and satellite cell protein-coding gene markers (*MYF5, PAX7*). The spatial distribution of C) *H19*, D) *MEG3*, E) *CARMN*, and F) *CASC15,* all with the same magnification, location, and protein-coding MERFISH probes as box inset in B).

*H19* is an imprinted lncRNA, and its pre-training abundance in human muscle is associated with training-induced changes in maximal aerobic capacity ^77^. Spatially, it was ubiquitous throughout human skeletal muscle (Figure 4C) without any strong associations with fibre type markers or mononuclear cell types (Figure 4A). In contrast, Maternally Expressed Gene 3 (*MEG3)*, another imprinted lncRNA, was co-localized with satellite cell gene markers (*PAX7* and *MYF5*; Figure 4D), and its expression was significantly correlated with *PAX7* expression (r =0.51; FDR = 2.48 x10^-29^; Figure 4A) in the bulk transcriptomic data. Indeed, a previous report in mice ^78^ found that *MEG3* is enriched in satellite cells and modulates myoblast differentiation and proliferation, impacting skeletal muscle regeneration.

Cardiac mesoderm enhancer-associated non-coding RNA (*CARMN*) had a low average expression in bulk human skeletal muscle (4.4 log_2_ units), implying it may not be expressed by mature muscle cells. Figure 4E demonstrates that *CARMN* appears located near endothelial cell gene markers (*APLNR, TIE1, KDR, ENG*), and we noted that it was positively associated with endothelial cell gene markers (e.g., *TIE1*: r =0.33; FDR = 3.26 x10^-12^; Figure 4A) in our bulk transcriptomic data. Previous work claimed that *CARMN* is a vascular smooth muscle cell-specific lncRNA ^79^. Although vascular smooth muscle cells are absent from muscle capillaries ^62^, pericytes are associated with endothelial cells in skeletal muscle and have been proposed to regulate endothelial cell proliferation and differentiation ^80^. Notably, in our data, *CARMN* expression is most strongly associated with pericyte gene markers (e.g., RGS5: r =0.50; FDR = 4.31 x10^-27^; Figure 4A), which leads us to hypothesize that *CARMN* might modulate the role of pericytes during skeletal muscle growth. Many reports of cell-unique patterns of lncRNA expression come from a failure to appreciate the limitations of RNAseq cDNA library production^81^. Like *CARMN* expression, cancer susceptibility candidate 15 (*CASC15*) also has a low signal in bulk skeletal muscle (Figure 4E). *CASC15* is an intergenic lncRNA, and small interfering knockdown experiments have shown that *CASC15* contains tumour-promoting properties in melanoma cell lines ^82^, and we find greater expression of *CASC15* is associated with human muscle growth *in vivo*.

### Data-driven RNA network modelling of ncRNA and protein-coding genes demonstrates that the hypertrophy-related ncRNAs represent functionally interacting pathways

Given most ncRNAs have no known function despite representing half the genome, inference methods are required to highlight potential molecular functions. We applied time-consuming data-driven quantitative network modelling to the muscle coding and noncoding transcriptome (n=437) and identified statistically robust co-expression modules that contained hypertrophy-related ncRNAs. There were 136 significant co-expression modules that contained at least one hypertrophy-related ncRNA (Figure 5A). Twelve of these modules were enriched in gene ontology biological processes, including cell migration, contractile fibre, inner mitochondrial membrane complex, RNA processing, and MHC protein complex processes (Table 1; Supplement Data S8 - 10). Application of a network-based enrichment method^68^ to the co-expression modules independently identified by MEGENA, in independent data, identifies that the hypertrophy ncRNA-containing modules interact at least two levels (Figure 5B). Firstly, genes belonging to modules contribute to more than one distinct biological function (pie-chart structures) both within and across functional groups (e.g. regulation of cell activation and positive regulation of cytokine production). Secondly, functional group labels highlight muscle tissue level processes (e.g. muscle structural development and vascular development) or the involvement of low abundance cells (e.g. lymphocyte activation). The genes driving these significant overlapping interactions are listed in Supplement Data S11.

**Figure 5.**
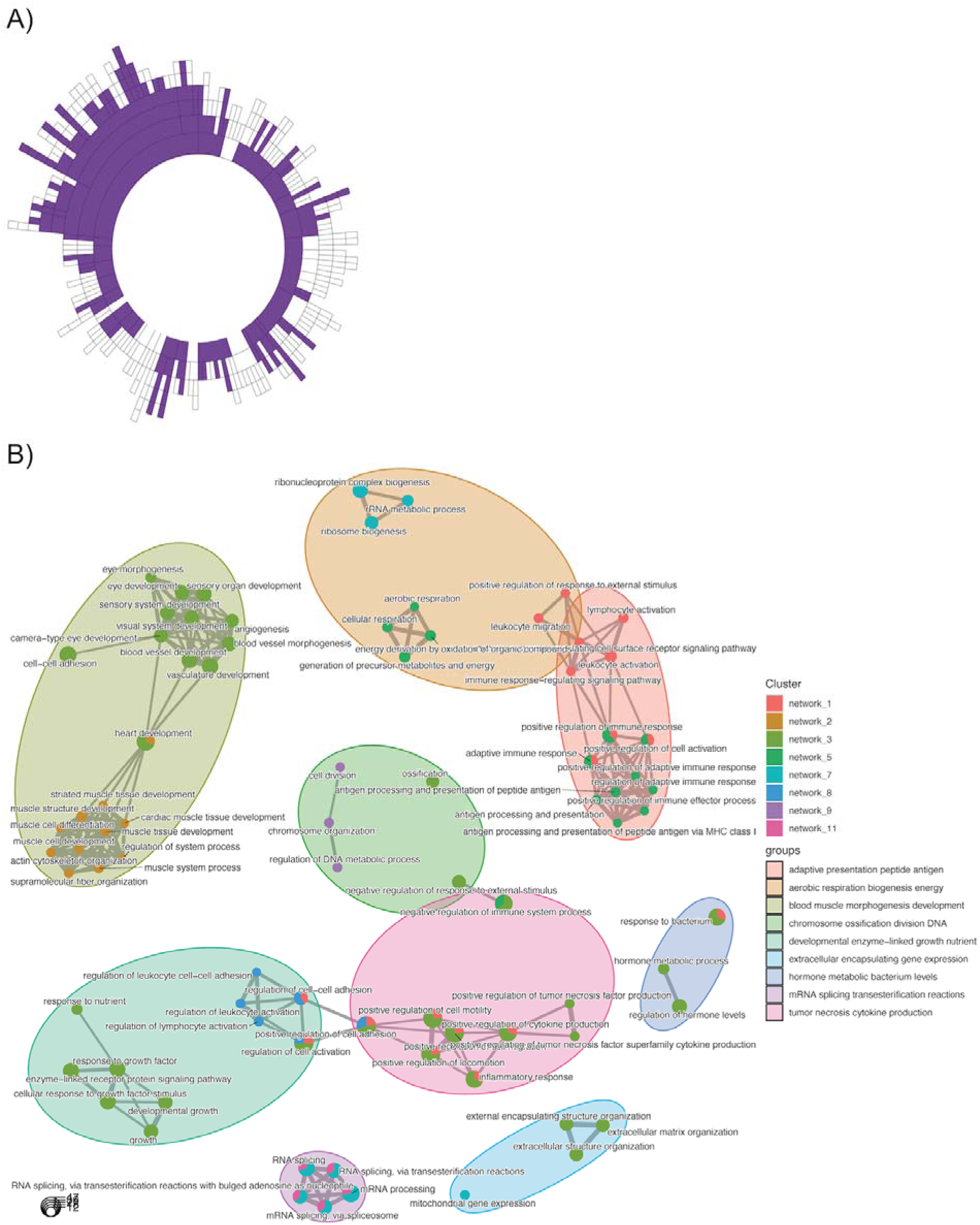
Discovery of interacting functional pathways that contain hypertrophy-related ncRNA genes. A) Sunburst plot denoting the hierarchical organisation of ncRNA containing co-expression network modules constructed from 437 resting skeletal muscle biopsy samples (< 50 years old), using MEGENA ^35^. Each concentric ring corresponds to a cluster of co-expressed genes, with the size of the ring being proportional to the number of genes. Purple concentric rings are modules that contain at least one of the 110 hypertrophy-related ncRNA genes. B) Using the compareCluster function^67,68^ in the ClusterProfiler R package and the GOTERM_BP_ALL from DAVID as the ontology database, and the ∼13.4K protein coding genes expressed in muscle, we established the relationships between hypertrophy ncRNA containing modules. A q-value of 1% was utilized as a filter for presentating significant ontologies. Pie-charts illustrate when genes from more than one network contribute to an individual significant ontology catergory, while some ontology catergories extend across distinct grouped functions. Further, each network can contain multiple ontologies which also show overlap within and across groups of functional processes. Finally, there is evidence that interacting groups of functions occur at the level of immune cell types. This analysis demonstrates that the ncRNAs interact at both the pathway and cell interaction level in a coordinated manner.

**Table 1.**
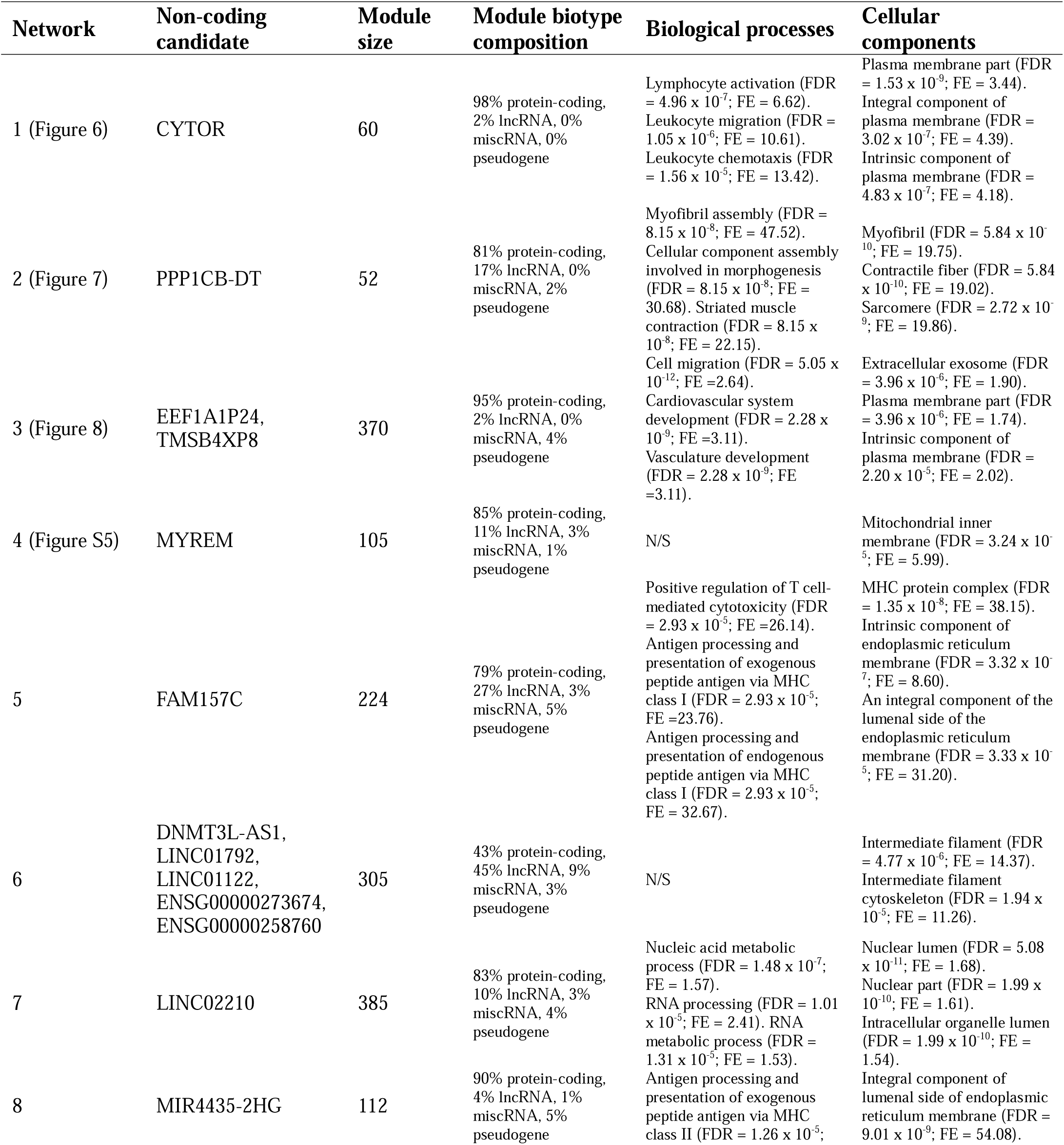
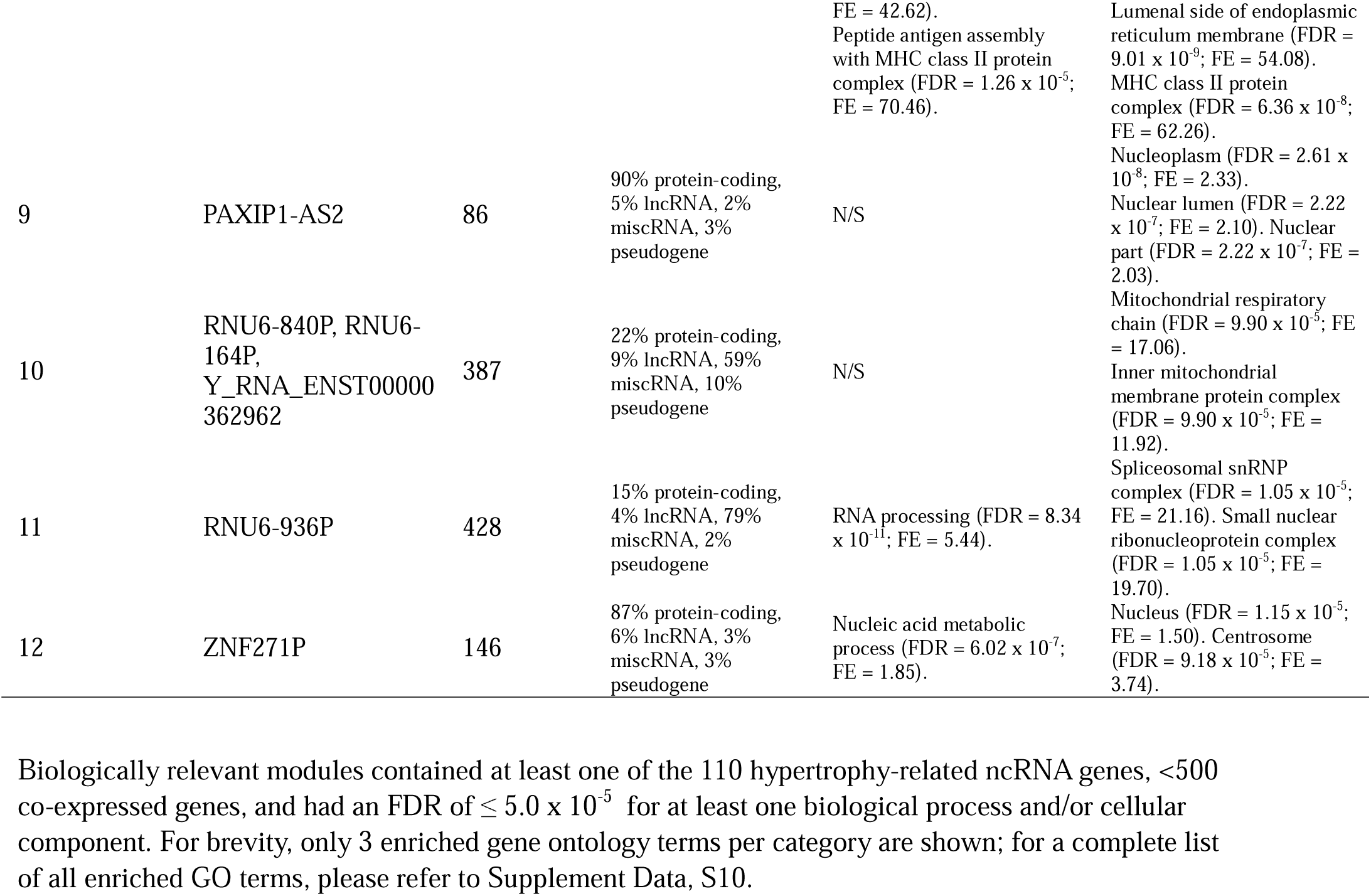
Summary of biologically relevant significant RNA co-expression modules containing hypertrophy-related ncRNA genes.

We next explored, in fine detail, some of the key MEGENA modules. One containing 60 genes included *CYTOR* – a ncRNA known to regulate hypertrophy in model systems - (Figure 6A; Table 1) – and the module represented immune-related processes e.g. leukocyte migration (FDR = 1.05 x 10^-6^; fold enrichment [FE] = 10.61), lymphocyte activation (FDR = 4 x 10^-7^; FE = 6.62), and leukocyte chemotaxis (FDR = 1.56 x 10^-5^; FE = 13.42). The hub gene of this module was lymphocyte cytosolic protein 1 (*LCP1*, or L-plastin), which has a direct role in aggregating actin filaments into parallel bundles, which is critical for T-cell activation, migration, and stability ^83^. *LCP1* expression was positively correlated with *CYTOR* expression (r = 0.49) in the bulk muscle transcriptome (Figure S3A). *FCER1G*, which encodes the Fc Epsilon Receptor Ig protein, facilitates the activation of natural killer cell lymphocytes, and it was also positively correlated with *CYTOR* (r = 0.66; Figure S4A). The median expression of the network genes in skeletal muscle is very low (3.7 log_2_ units; Figure 6B), indicating that this module of co-expressed genes is likely expressed in the small number of immune cells found in muscle tissue – illustrating the sensitivity of our transcriptomic methods. To further investigate, we reprocessed publicly available data from the same laboratory platform, obtained from human primary immune cells (macrophages [GSE124106], CD8+ T cells [GSE117230], and CD4+ T cells [GSE62921], endothelial cells [GSE236137], and our own *in vitro* human myotube data ^27^. We noted that *LCP1*, *FCER1G*, and *SLC31A2* are mostly expressed in immune cells (macrophages, CD8+ and/or CD4+ T-cells), compared with other cell types found in skeletal muscle tissue samples (Figure 6C). This trend is observed for most genes in the *LCP1* module (Figure S4B). Thus, *CYTOR* regulation is linked to successful muscle hypertrophy in humans and is found in an immune-cell-related module.

**Figure 6.**
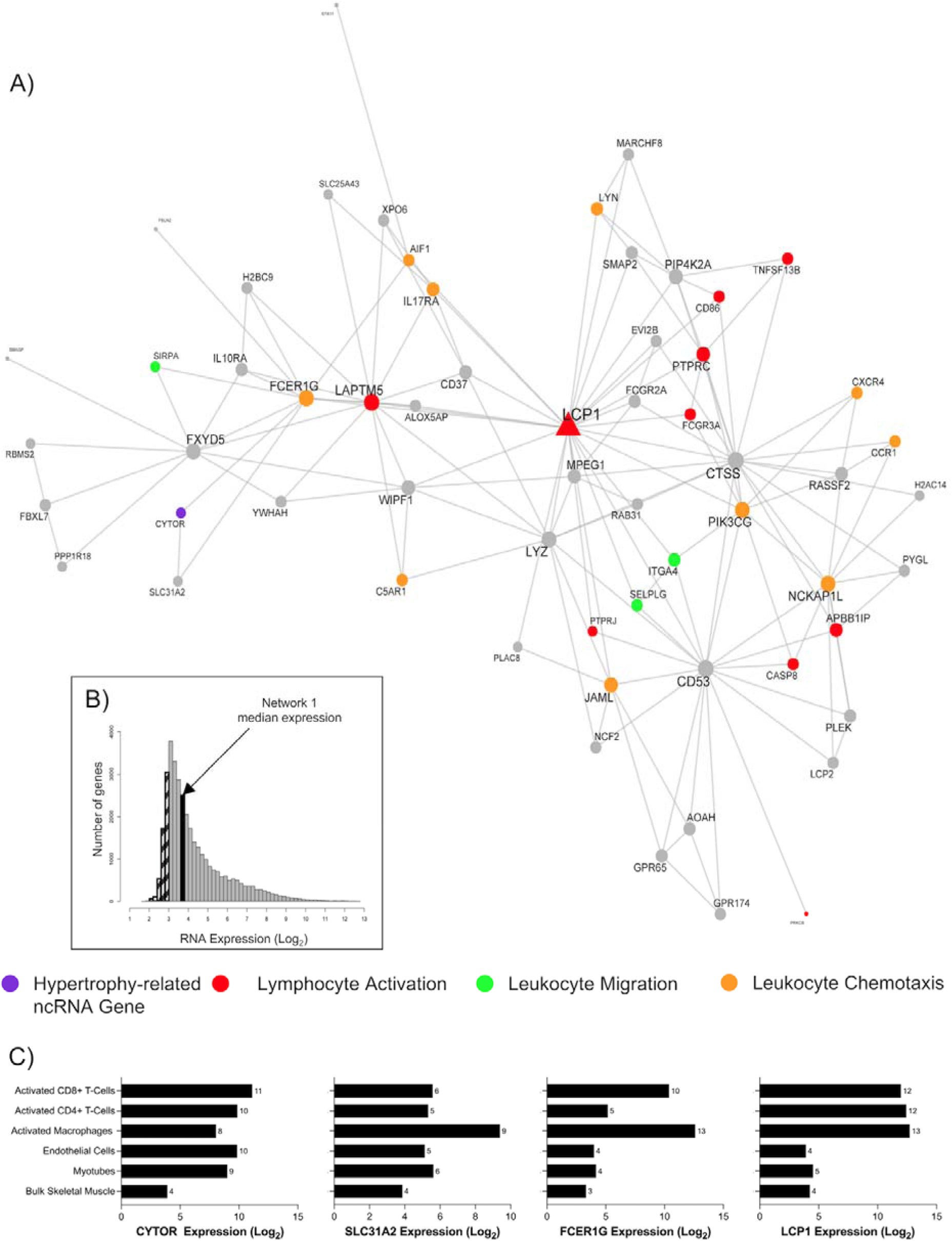
*CYTOR* is the only ncRNA gene co-expressed in network 1 and biologically associated with immune cell-related functions. A) A biologically enriched gene co-expression network (network 1; table 1), which contains the hypertrophy-related ncRNA gene, *CYTOR* (purple). This network of genes is related to immunological biological processes, such as lymphocyte activation (red), leukocyte migration (green), and leukocyte chemotaxis (orange). Node and label size are proportional to node degree value within each network, and triangle symbols denote genes that are hubs of the cluster of co-expressed genes. B) A frequency plot of the median log_2_ expression of 35k genes in resting bulk skeletal muscle. Hatched bars reflect gene signals most likely indistinguishable from background signal ^15^. The black bar denotes the median log_2_ expression of all 60 genes apart from network 1. C) The log2 expression of several network 1 genes (*CYTOR*, *SLC31A2*, *FCER1G*, *LCP1*) in bulk skeletal muscle, human myotubes, endothelial cells, activated macrophages, CD8+ T cells, and CD4+ T cells.

The next module investigated (Figure 7A; Table 1) centred around the slow skeletal muscle troponin isoform, *TNNT1*. This module was composed of genes robustly co-expressed in muscle cells, median expression of 6.4 log_2_ units (Figure 7 and Figure S5). Nested within this module was a hypertrophy-related ncRNA gene, phosphatase 1 catalytic subunit beta divergent transcript (*PPP1CB-DT*; Figure 7A). Based on DAVID analyses, the module was enriched in biological processes related to myofibril assembly (FDR = 8.15 x 10^-8^; FE = 47.52), striated muscle contraction (FDR = 8.15 x 10^-8^; FE = 47.52), muscle filament sliding (FDR = 8.15 x 10^-8^; FE = 22.5), and the ontology label ‘transition between fast and slow fibre’ (FDR = 2.80 x 10^-7^; FE = 217.81) above the muscle transcriptome. Attractin-like-1 (*ATRNL1*) is in this module, and we previously identified ^25^ *ATRNL1* was correlated with human skeletal muscle growth in a targeted analysis. The module also contained estrogen-related receptor gamma (*ESRRG*), which contributes to exercise-mediated remodelling of skeletal muscle function^84,85^. In mouse models, *ESRRG* promotes an increased number of oxidative muscle fibres and is considered a regulator of oxidative metabolism ^86^. The expression levels of most genes in this module are positively associated with markers of type I fibres and negatively associated with markers of type II fibres (Figure S5C). Collectively, the *TNNT1* module most likely reflects genes that enhance the oxidative capacity of skeletal muscle, a common occurrence with RT in humans ^87,88^.

**Figure 7.**
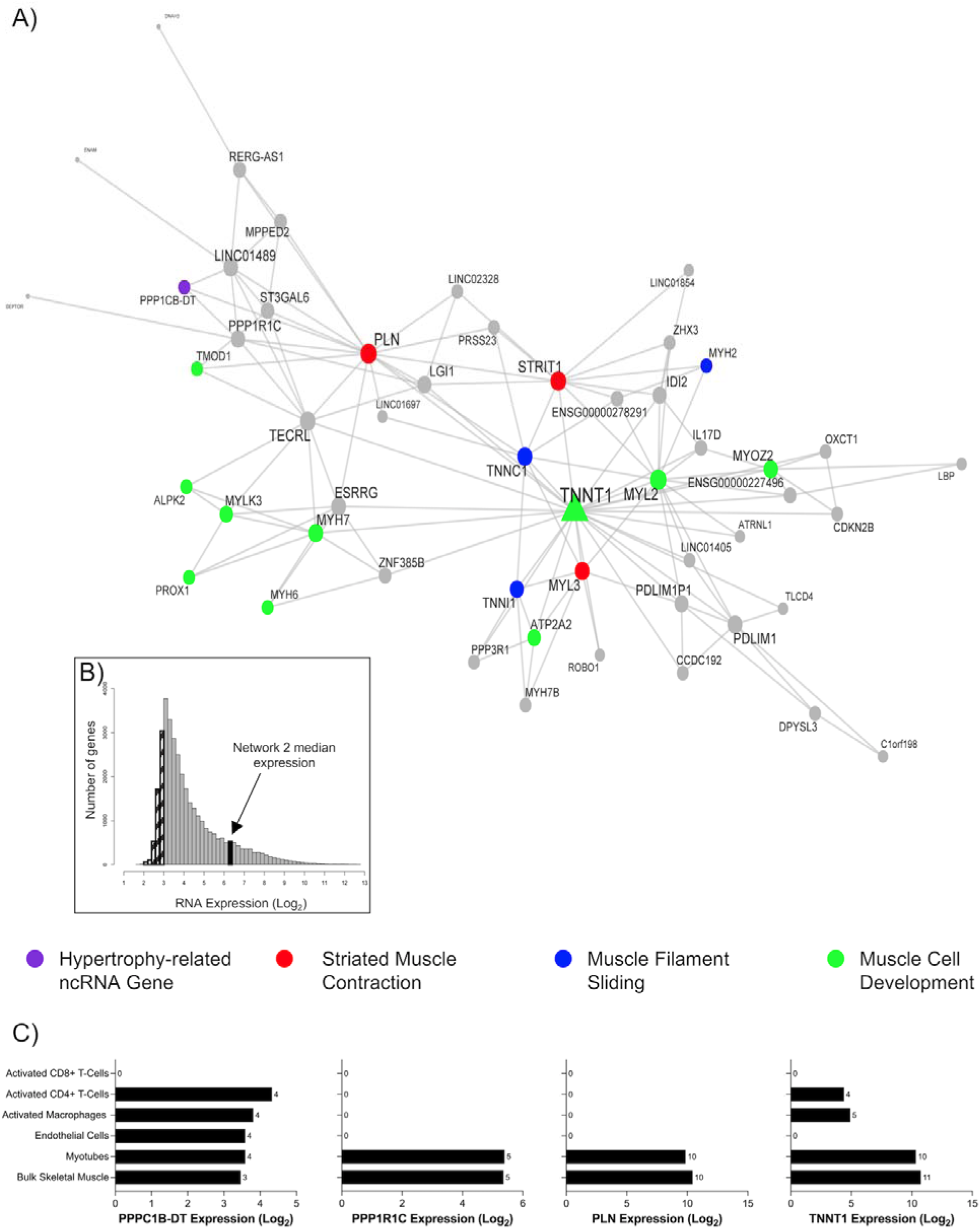
A highly expressed gene co-expression network in skeletal muscle is related to muscle remodelling processes. A) Network 2 is composed of 52 genes and contains the hypertrophy-related ncRNA gene, phosphatase 1 catalytic subunit beta divergent transcript (*PPP1CB-DT*; purple). This network of genes is related to skeletal muscle remodelling processes, such as striated muscle contraction (red), muscle filament sliding (blue), and muscle cell development (green). Node and label size are proportional to node degree value within each network, and triangle symbols denote genes that are hubs of the cluster of co-expressed genes. B) A frequency plot of the median log_2_ expression of 35k genes in resting bulk skeletal muscle. Hatched bars reflect gene signals most likely indistinguishable from background signal ^15^. The black bar denotes the median log_2_ expression of all 52 genes apart from network 2. C) The log2 expression of several network 2 genes (*PPPC1B-DT*, *PP1R1C*, *PLN*, *TNNT1*) in bulk skeletal muscle, human myotubes, endothelial cells, activated macrophages, CD8+ T cells, and CD4+ T cells.

A final module we considered in detail (370 genes; Figure 8; Table 1; Supplement Data, S9-10) was centred around *MAN1A1* and the *ATLASTIN GTPase 3 (ATL3)* genes, the latter of which are a constitutive endoplasmic reticulum fusion protein^89^. Loss of function mutations in *ATL3* lead to muscle weakness^90^. This large module was enriched in GO terms related to microvascular remodeling e.g., angiogenesis (FDR = 2.77 x 10^-5^; FE = 3.55), blood vessel development (FDR = 6.36 x 10^-8^; FE = 2.97), regulation of vasculature development (FDR = 1.66 x 10^-6^; FE = 3.71), and circulatory system development (FDR = 1.66 x 10^-8^; FE = 2.55). Network-based enrichment analysis^68^ identified that these processes were directly linked to the muscle remodelling pathways. *EEF1A1P24* and *TMSB4XP8* are two hypertrophy-related ncRNA genes in this module, and both are pseudogenes (Figure 8; purple). Interestingly, they are part of a correlative cluster of several other pseudogenes (median R = 0.60; *EEF1A1P9, EEF1A1P17, EEF1A1P25, EEF1A1P22, EEF1A1P16, EEF1A1P6, TMSB4XP4, TMSB4XP2*; Figure 8 box inset). Pseudogenes are defined as copies of parent genes that are either retro-transposed counterparts or processed mRNAs (i.e., processed pseudogenes) or harbour mutations derived from gene duplication (i.e., unprocessed pseudogenes) ^91^. In recent years, these genes have been identified as potential decoys for microRNAs and modulators of the sensitivity of the related protein-coding genes to inhibition ^92^.

**Figure 8.**
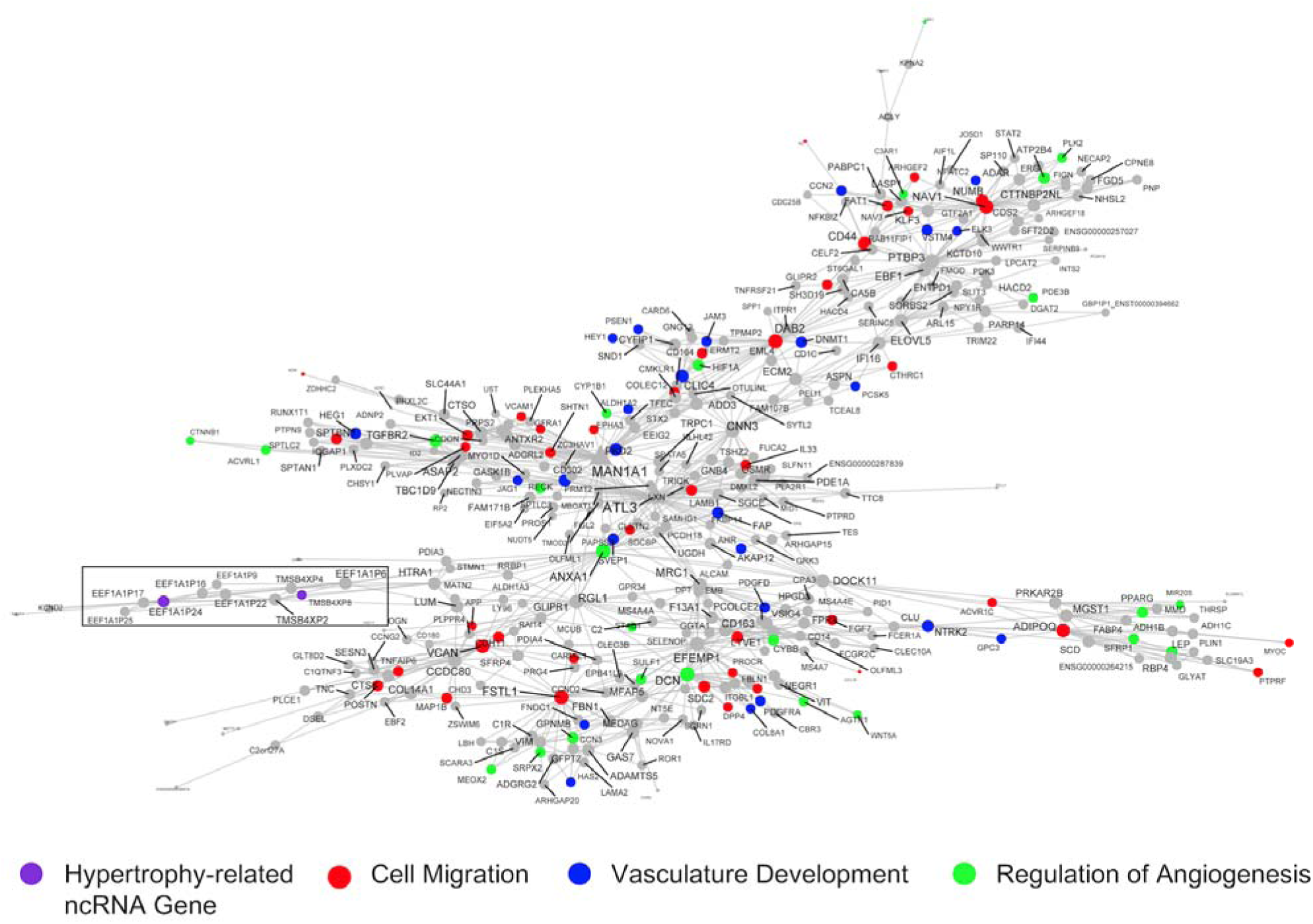
RNA network related to vascular remodelling harbours a pair of pseudogenes related to skeletal muscle hypertrophy. This network is one of the largest gene co-expression RNA networks (370 genes), and a significant proportion of genes are associated with cell migration (red), vasculature development (blue), and regulation of angiogenesis (green). Embedded in this network are a cluster of pseudogenes (*EEF1A1P9, EEF1A1P24, EEF1A1P17, EEF1A1P25, EEF1A1P22, EEF1A1P16, EEF1A1P6, TMSB4XP4, TMSB4XP2, TMSB4XP8;* black box), two of which are hypertrophy-related ncRNA genes (*TMSB4XP8* and *EEF1A1P24,* purple). Node and label size are proportional to node degree value within each network, and triangle symbols denote genes that are hubs of the cluster of co-expressed genes.

## DISCUSSION

The present study is a comprehensive profile of the human skeletal muscle ncRNA transcriptome, which successfully identified numerous hypertrophy ncRNA candidates embedded in tissue growth pathways, often in a cell-specific manner and that they represent modules of integrated overlapping pathways. Most of the 110 ncRNA genes linked to successful hypertrophy would not have been identified had we relied on a typical RNA sequencing workflow. The use of data-driven quantitative RNA network modelling in independent data, along with network-based pathway enrichment and single-cell spatial transcript profiles allowed us to identify the functional pathways and cell types these ncRNA genes operated within. We established that type II myofibril-associated *MYREM* demonstrates specific myonuclear domain expression *in vivo* in human muscle. This work also added to the growing body of evidence that local immune-related responses, mitochondrial remodelling, and angiogenesis are all critically important contributors to successful human skeletal muscle hypertrophy ^23,93,94^.

### The benefits of high-density array profiling to study the skeletal muscle ncRNA transcriptome

A majority of recent skeletal muscle transcriptome studies have relied on short-read RNA sequencing ^28,71,72,95–98^. We have shown that short-read RNA sequencing profiling of human skeletal muscle detects only ∼20% of constitutively expressed ncRNAs and demonstrates enormous inter-study heterogeneity^71,72^ (See Figure 3E). In the present study we estimate that >7000 ncRNA genes are detected in every skeletal muscle sample (Figure 3A), consistent with our previous work^13,15,27,46^. Notably, only 30% of the 110 hypertrophy ncRNA genes were consistently detected in two short-read RNA sequencing studies of skeletal muscle growth (Figure 3E). Stochastic properties of sequencing and the reliance on poly-A enrichment strategies (ncRNAs are deficient in poly-A tails) means that most ncRNAs are not detected. A striking example is *H19*, a hypertrophy and exercise-related ncRNA that is abundantly expressed in human skeletal muscle (Supplement Data, S8), yet a recent RNA sequencing study indicated that H19 was not ‘robustly expressed’ in human skeletal muscle ^28^. In 2012, we identified *H19* as being part of a baseline gene expression signature predictive of the cardiovascular adaptive response to exercise training ^77^. More recent work demonstrated that the *H19* locus encodes several miRNAs that regulate protein translation initiation factors implicated in protein synthesis ^99^. Interestingly, *H19* directly interacts with the C-terminal domain of dystrophin, blocking its degradation^100^. Dystrophin links the internal cytoskeleton to the extracellular matrix ^101^, and transcriptome responses to exercise are linked to Dystrophin status ^69^. In Duchenne Muscular Dystrophy (DMD), *H19* interaction with dystrophin is impaired, resulting in the degradation of dystrophin and muscle degeneration ^100^. Clearly, all of this evidence indicates that *H19* is a crucially important lncRNA in muscle (with wide-spread expression; Figure 4C) and directly contributes to regulating tissue remodelling.

### Identifying the spatial localisation of ncRNA genes aids in generating functional hypotheses

Skeletal muscle is a conglomeration of cells dominated by large multinucleated myofibers while containing mononucleated cells, including satellite cells, endothelial cells, immune cells, and pericytes. There is growing support that these rarer cells contribute to skeletal muscle remodelling and that perturbation of their role is related to muscle dysfunction ^58,102,103^. Using merfish probes, we illustrated that our hypertrophy ncRNA *MYREM* was localised near the periphery of skeletal muscle fibres and co-stained with DAPI indicative of mature myonuclei ^104^. Previous *in vitro* work showed that *MYREM* was markedly abundant in the nuclei of myotubes compared with the nuclei of myoblasts ^37^. Our data indicate that *MYREM* appears to be a novel marker of human myonuclei *in vivo* (absent from cells expressing endothelial or satellite cell marker genes). Nuclear lncRNAs participate in vital processes, such as chromatin organisation, transcriptional regulation, and RNA processing ^105^. *MYREM* interacts with the RNA-binding protein (RBP), heterogeneous nuclear ribonucleoprotein L (HNRNPL) to enhance the transcription of the fast-twitch isoform skeletal muscle myosin-binding protein C2 (*MYBPC2*). *MYBPC2* is involved during the later stages of the myogenic program, whereby mononucleated myoblasts fuse into multinucleated myotubes, and the myosin heavy chain proteins are expressed and function in muscle contraction ^106^. Interestingly, network modelling indicated that *MYREM* is co-expressed with a mitochondrial-inner membrane gene expression module (Figure S6) containing genes such as *ATP5F1D*, *NDUFA3*, but also the protein-translation initiation factor *EIF3K* ^107^. Remodelling of the mitochondria during hypertrophy-induced myogenesis may support increased energetic-related mechanisms (e.g., translation^108^), explaining why a myogenesis lncRNA is co-expressed with mitochondrial-related genes and given this may mostly reflect the physiological loading of type II muscle fibres.

To our knowledge, we are the first to report that *MEG3* localises rather selectively to muscle satellite cells in human skeletal muscle (Figure 4D), while *MEG3* has been reported to be nuclear located in other cell types ^37^. *MEG3* is a maternally imprinted gene known to have tumour-suppressive properties in various cancers ^109^. Interestingly, *Gtl2*, the mouse ortholog of *MEG3* ^110^, targets the polycomb repressive complex 2 to epigenetically silence a variety of genes from the *Dlk1-Dio3* locus, including the protein-coding transmembrane gene, *Dlk1* ^111^. A muscle-specific *dlk1* knockout in mice has altered satellite cell lineage progression following skeletal muscle injury ^112^, and overexpression of *dlk1* is associated with pathological hypertrophy ^113,114^. Taken together, the spatial location, network and whole tissue bulk correlation analysis indicates that several hypertrophy-related ncRNA genes in skeletal muscle are likely mediating their influence via rare cell types (by abundance) in skeletal muscle.

### Data-Driven Network Analysis Provides Biological Insight into Novel ncRNAs

Most of the hypertrophy-related ncRNAs in the present study had no previous roles in skeletal muscle biology (Table S2). For example, *PPP1CB-DT* was co-expressed in a module of genes related to contractile fibre remodelling (Figure 7). *PPP1CB-DT* is a divergent ncRNA to *PPP1CB*, a protein-coding gene for one of three catalytic subunits of protein phosphatase 1 (*PP1*), which is a serine/threonine-specific protein phosphatase that plays major roles in regulating skeletal muscle contractility ^115^. Divergent ncRNAs are transcribed in the opposite direction from the promoters of protein-coding genes ^116^ and may affect adjacent protein-coding genes through direct regulation in *cis* ^117^ or by participating in chromatin modifications ^118^. We also noted that a pair of hypertrophy-related pseudogenes, *EEF1A1P24* and *TMSB4XP8*, were found to be regulated in a vascular remodelling and angiogenesis module (Figure 8). The angiogenic molecular program is tightly coupled to skeletal muscle growth ^25,119^, and enhancing angiogenesis (by aerobic conditioning) may augment growth ^120^. *TMSB4XP8* is a thymosin beta 4-X (*TMSB4X*) pseudogene, an actin-binding protein with known roles in cell migration and angiogenesis ^121^. In addition, *EEF1A1P24* is a processed pseudogene for the protein-coding gene, eukaryotic elongation factor 1 (*EEF1A*), which plays an important role in delivering the amino-acyl tRNA to the ribosome during protein synthesis ^122^. Until recently a broad misunderstanding is that pseudogenes are non-functional counterparts to their related genes ^91^. Pseudogenes exert biological functions in a DNA (facilitating chromatin structure remodelling or gene conversion ^91^), RNA (competitive endogenous RNA networks ^123^), and/or via protein (some pseudogenes contain protein-coding potential ^6^) dependent manner. We demonstrated, using network-based pathway modelling on large-scale quantitative network models of independent muscle transcriptomes (n=437), that a group defined as ‘blood muscle morphogenesis and development’ directly connected the angiogenesis MEGENA modules with muscle tissue development (Figure 5B). These vascular-related ncRNAs may aid modification of microvascular flow during exercise ^124^ or feeding^125^, which helps contribute to meeting the demands of increased rates of protein synthesis and overall a key role for the vascular system modulating muscle hypertrophy is now apparent ^25,119,120^.

### RNA networks reveal novel roles of ncRNA genes in skeletal muscle immunomodulation

The immune system also appears to be a novel regulator of skeletal muscle remodelling^103^. Our ncRNA analyses highlight biological processes distinct from our previously described protein-coding hypertrophy signature ^25^, such as immune-cell-related functions (Figure S7). For example, *MIR4435-2HG* (a miRNA host gene) was co-expressed in a module dominated by immune-related processes (e.g., peptide antigen assembly with MHC class II protein complex; Table 1; Supplement Data, S10), and *MIR4435-2HG* has most recently described as a gene called ‘*Morrbid’*. *Morrbid* tightly regulates the lifespan and turnover of myeloid cells (e.g., neutrophils and eosinophils) by regulating the transcription of pro-apoptotic genes ^95,126^. Notably, *CYTOR* was also co-expressed in an immune-dominated protein-coding RNA module (Figure 6), with robust expression in macrophages, CD8+ T cells, and CD4+ T cells (Figure S4B). To date, the characterisation of *CYTOR* in mouse muscle tissue has focused on muscle cells and not other cell types found within muscle tissue ^95^. In muscle cells, *CYTOR* sequesters the *TEAD1* transcription factor ^95^, which can transcriptionally regulate the small peptide apelin that regulates paracrine communication between muscle cells and endothelial cells ^127^. Apelin is a feature of the protein-coding pathways associated with human muscle growth. We find that *CYTOR* was widely expressed across multiple cell types (Figure 6C), suggesting that *CYTOR* can impact skeletal muscle hypertrophy via multiple mechanisms. The present analysis suggests a model whereby a number of rare but important resident secretory cell types may regulate adaptation and release factors in a paracrine manner (e.g. a non-muscle source for various proposed myokines) to promote skeletal muscle remodelling ^103^. Collectively, our analyses reveal that low-abundance cell types can be directly studied in bulk transcriptomic profiles of human skeletal muscle tissue.

## LIMITATIONS

The present human clinical studies are not without its limitations. Four out of the five exercise loading studies relied on DXA to quantify changes in LLM, and the hydration status of the participant can influence this measure and contribute to the clinical variation^128^ and this may have reduced our ability to identify ncRNA hypertrophy candidates. With respect to the interpretation of our results, although ncRNAs are predicted to contain no protein-coding potential, some ncRNAs contain small open reading frames that encode peptides ^6,12^. Without additional experiments, some of the hypertrophy-related ncRNAs identified in the current study may also reflect the actions of novel bioactive peptides. Nonetheless, detailed mechanistic studies will be required to establish the direct involvement of each ncRNA in the hypertrophic response. Nevertheless, the few ncRNAs known to have causal links to cell growth and muscle tissue hypertrophy were identified by our methodologies.

## ACKNOWLEDGEMENTS

JCM was supported by a Queens University Vice Principal Research Postdoctoral Fund. This work was also supported by Medical Research Council, UK through grants G1100015 and MR/Y010329/1 (JAT) and by the National Institutes of Health, USA (1R56AG061911-01 (CW and JT). This work was also funded by a grant to SMP from the National Science and Engineering Research Council (NSERC) of Canada. SMP is supported by the Canada Research Chairs Program.

## SUPPLEMENTAL MATERIALS

**Table S1.** Demographics of the five studies used for establishing ncRNA genes associated with skeletal muscle hypertrophy.

|  | Morton et al., 2019 <sup>40</sup> | Morton et al., 2016 <sup>39</sup> | Phillips et al., 2017 <sup>41</sup> | Mitchell et al., 2014 <sup>42</sup> | Stokes et al., 2020 <sup>25</sup> |
| --- | --- | --- | --- | --- | --- |
| Sample size, n | 32 | 33 | 47 | 20 | 12 |
| Age, years | 22 ± 3 (19 – 28) | 23 ± 3 (20 – 29) | 38 ± 9 (21 – 51) | 24 ± 3 (20 – 30) | 21 ± 3 (18 – 29) |
| Gender, M/F | 32/0 | 33/0 | 18/29 | 20/0 | 12/0 |
| Body mass index, kg/m<sup>2</sup> | 25 ± 6 (18 – 38) | 26 ± 9 (41) | 32 ± 4 (26 – 43) | 24 ± 4 (15 – 32) | 24 ± 3 (20 – 31) |
| Measurement instrument | DXA | DXA | DXA | MRI | DXA |
| Limbs Involved in Measurement | 1 | 2 | 1 | 1 | 1 |
| Pre-training LLM, kg | 10.3 ± 2.1 (7.2 – 14.6) | 24.2 ± 3.0 (17.4 – 30.9) | 5.8 ± 1.4 (3.2 – 8.6) | - | 9.5 ± 1.6 (7.7 – 13.4) |
| Post-training LLM, kg | 10.6 ± 2.1 (7.6 – 15.2) | 25.0 ± 3.0 (19.9 – 31.2) | 6.0 ± 1.4 (3.5 – 8.7) | - | 9.9 ± 1.6 (8.1 – 13.4) |
| Pre-training QMV, cm<sup>3</sup> | - | - | - | 1862.0 ± 402.6 (1039.0 – 2822.0) | - |
| Post-training QMV, cm<sup>3</sup> | - | - | - | 1985.0 ± 417.6 (1278.0 – 3104.0) | - |
| dLLM, % | 3.2 ± 3.1 (-3.7 – 9.1) | 3.0 ± 5.0 (-5.0 – 19.0) | 3.1 ± 3.7 (-4.5 – 11.0) | - | 5.0 ± 4.3 (-1.2 – 14.0) |
| dQMV, % | - | - | - | 7.1 ± 7.0 (-1.8 – 24.7) | - |
| R<sup>2</sup> dLLM (kg) vs Pre-training LLM (kg) | 0.07 | 0.01 | < 0.01 | - | < 0.01 |
| R<sup>2</sup> dQMV (cm<sup>3</sup>) vs Pre-training QMV (cm<sup>3</sup>) | - | - | - | < 0.01 | - |
| R<sup>2</sup> dLLM (kg) vs Age (years) | 0.26 | < 0.01 | < 0.01 | - | 0.04 |
| R<sup>2</sup> dQMV (cm<sup>3</sup>) vs Age (years) | - | - | - | 0.01 | - |
| R<sup>2</sup> dLLM (kg) vs Gender | - | - | 0.04 | - | - |
| LMR group, n | 20 | 20 | 24 | 14 | 10 |
| NMLR group, n | 10 | 13 | 20 | 5 | 2 |

Age, body mass index, Pre-training LLM, Post-training LLM, dLLM, Pre-training QMV, Post-training QMV, and dQMV are displayed as mean ± SD (min - max). Sample size, Gender, LMR group, and NMLR group are displayed as counts. Abbreviations: DXA, dual X-ray absorptiometry; MRI, magnetic resonance imaging; dLLM, delta leg lean mass; QMV, quadriceps muscle volume; LMR, lean mass responders; NMLR, no measurable lean mass response.

**Figure S1.**
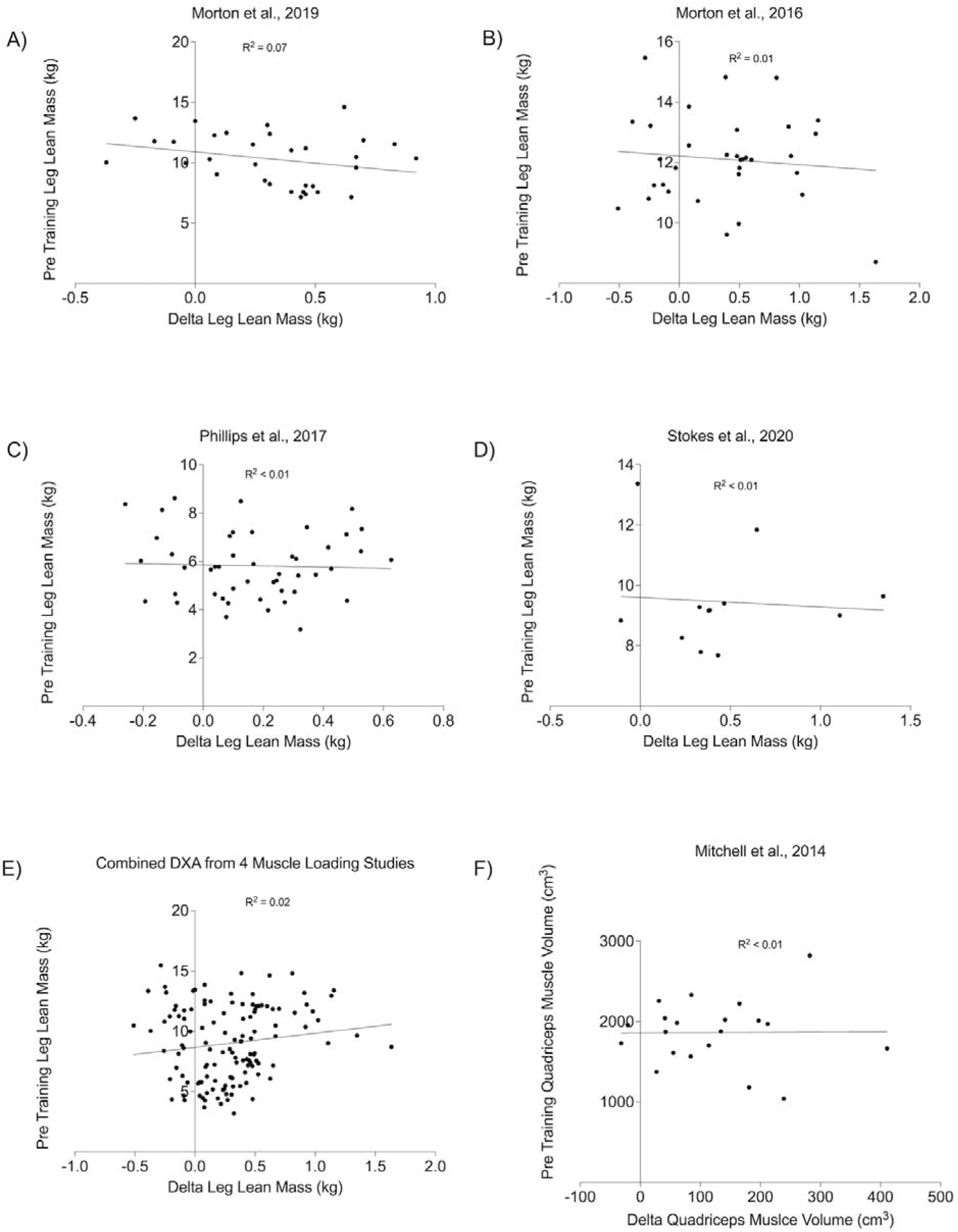
(A-D) Changes in leg lean mass versus pretraining leg lean mass for 4 muscle loading studies, and (E) depicts the aggregated relationship. F) Changes in quadriceps muscle volume vs pre-training quadriceps muscle volume.

**Figure S2.**
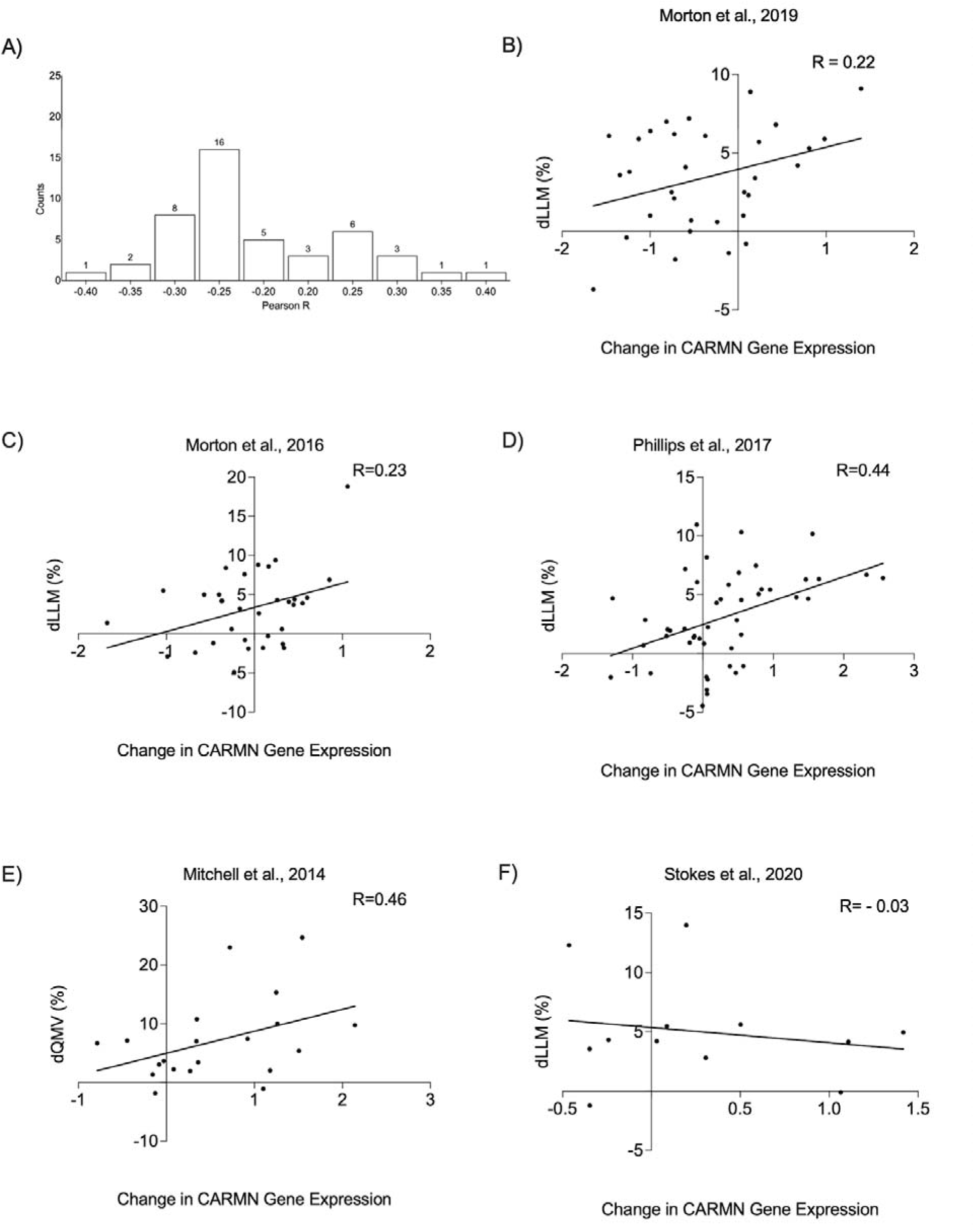
(A) Distribution of Pearson correlation coefficients among the 46 ncRNA genes containing a change in expression that was modestly associated with changes in LLM (dLLM), or changes in Quadriceps muscle volume (dQMV; Supplement Data S5). (B -F) Visual example from the linear modelling analysis, depicting the relationship between dLLM (or dQMV [E]) and changes in *CARMN* gene expression for each of the 5 individual exercise studies.

**Figure S3.**
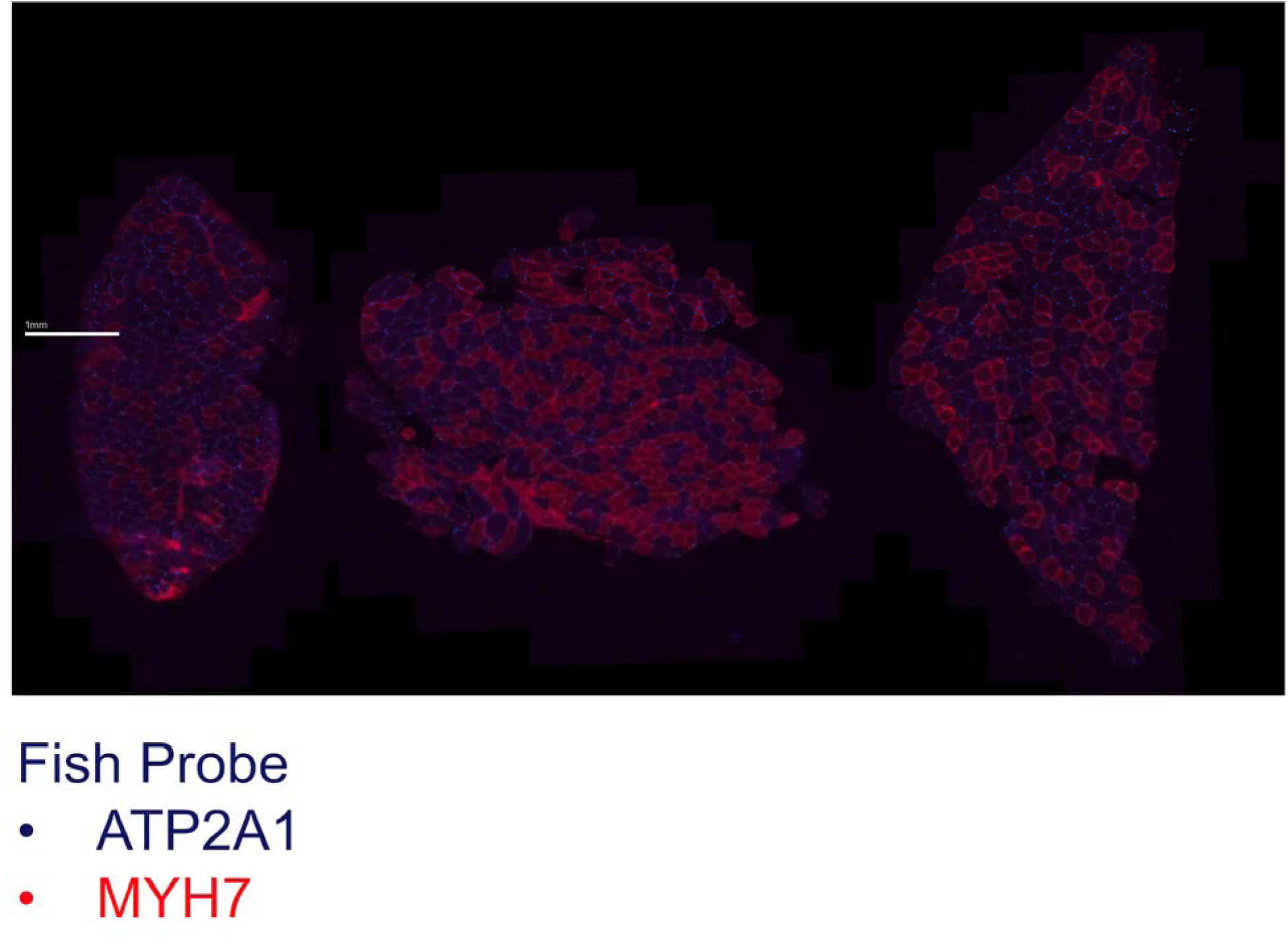
MERSCOPE-MERFISH overview for 3 samples used.

**Table S2.**
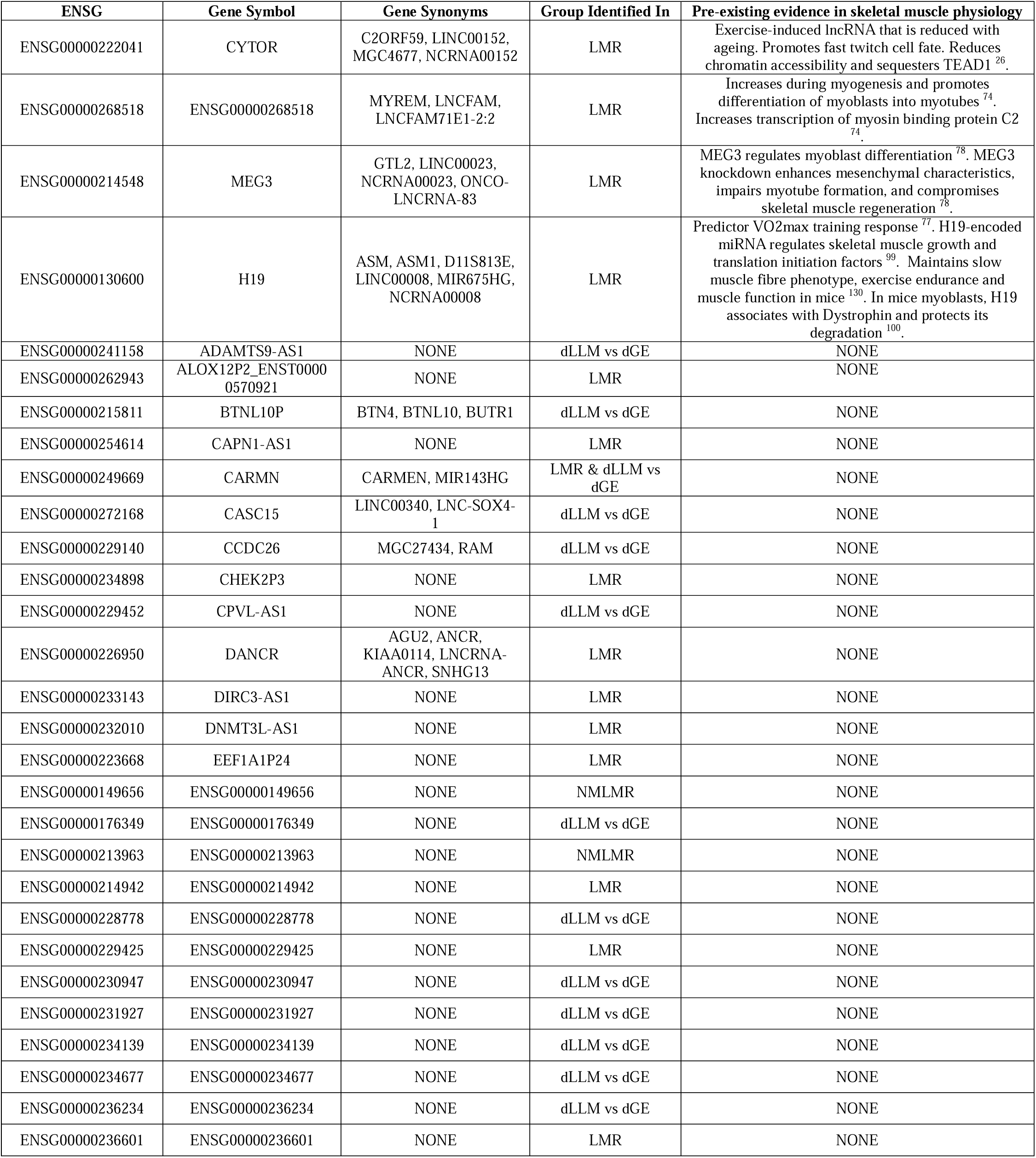
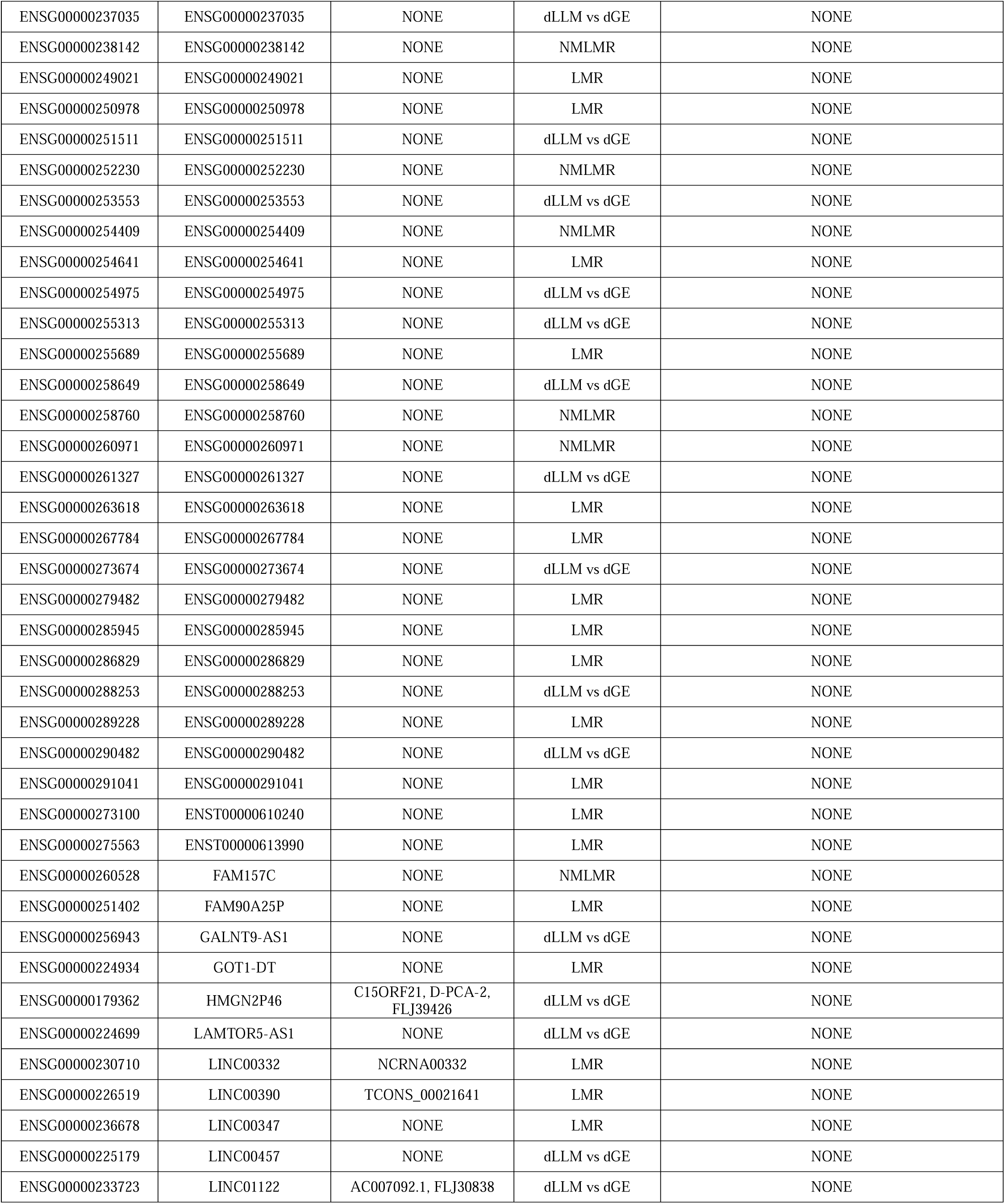
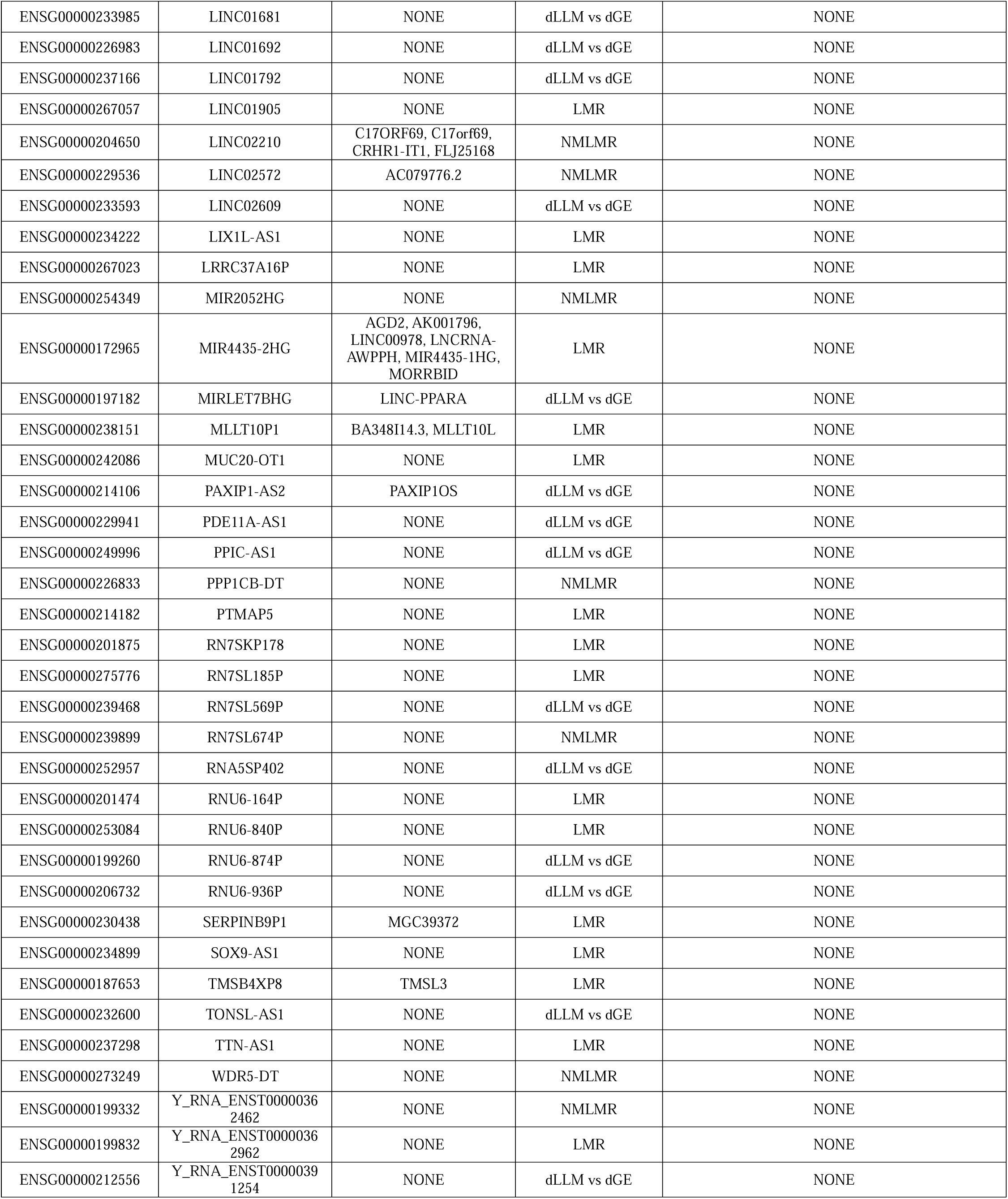
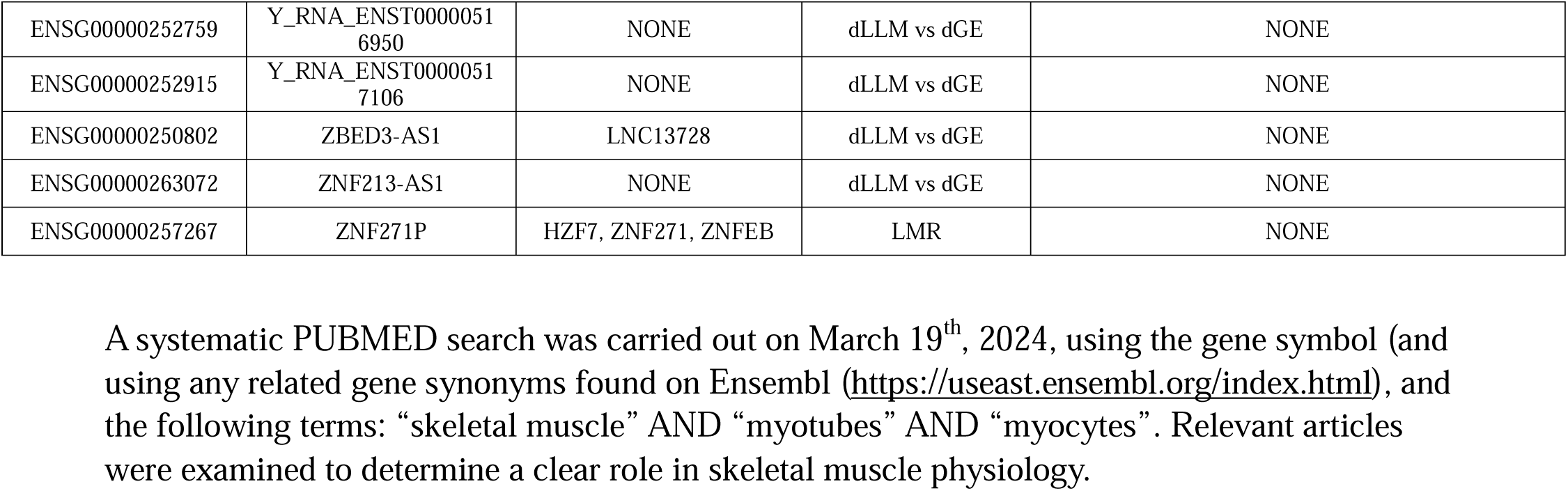
Existing biochemical relationship for 110 hypertrophy-related ncRNA genes in skeletal muscle.

**Figure S4.**
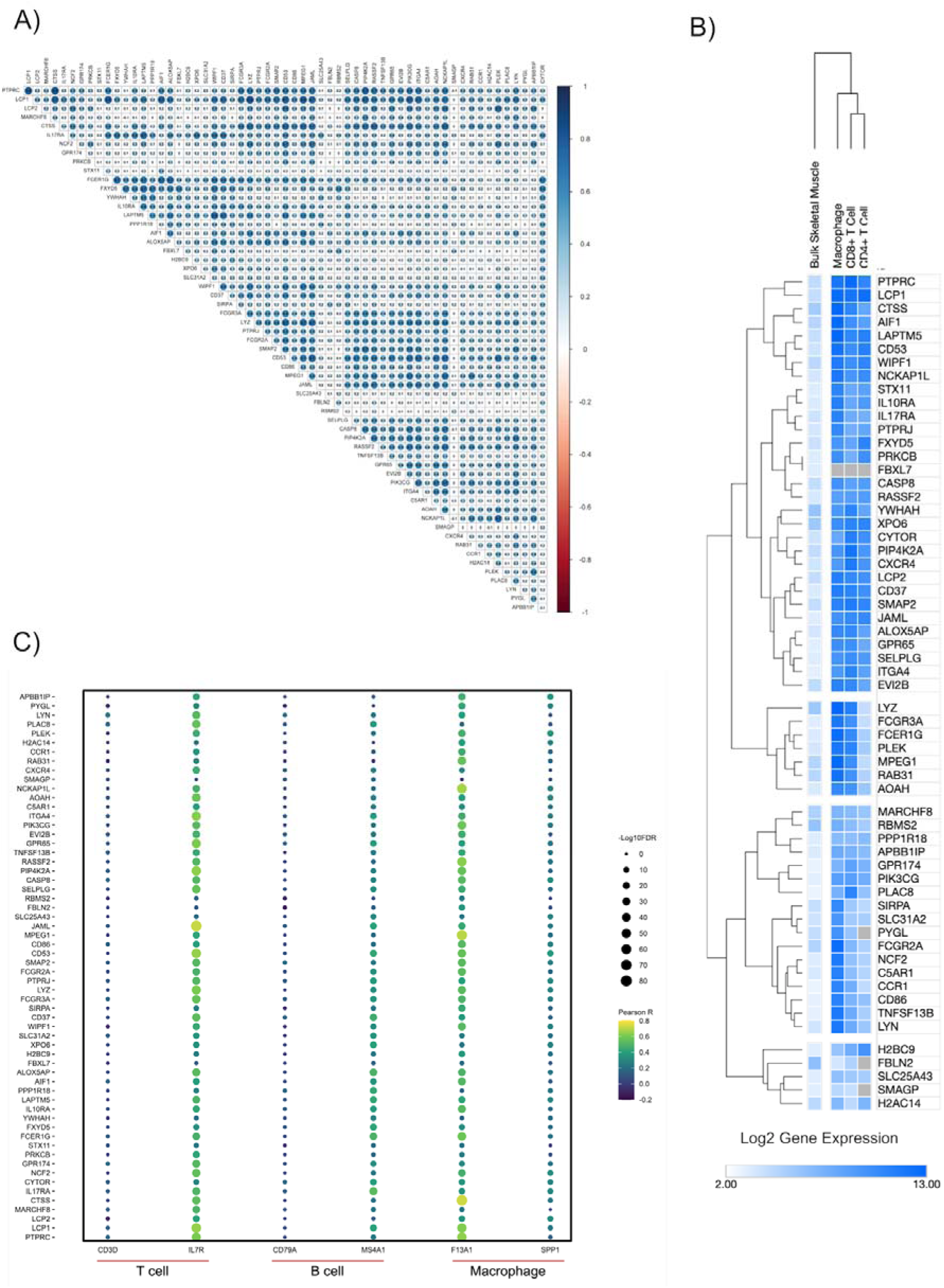
A) Pearson correlation matrix of all 60 genes co-expressed in network 1. Majority of the genes are positively correlated with each other in this network. B) the heatmap uses the log2 gene expression in skeletal muscle (n=437) and plots the 60 genes co-expressed in network 1 along with marker genes from three mononuclear cells of the immune system (macrophages, CD4 T-cells, and CD8 T-cells). The plot was created using Morpheus (https://clue.io/Morpheus), and genes and tissue types were hierarchically clustered using Euclidean distance (linkage method: complete). C) Dot plot depicting the association between the expression of network 1 genes, and gene markers for T-cells, B-cells and macrophages, in human skeletal muscle transcriptomic data (n=437). The colouring of the dot corresponds to the Pearson correlation coefficient, and the size of the dot is proportional to the –Log^10^ FDR.

**Figure S5.**
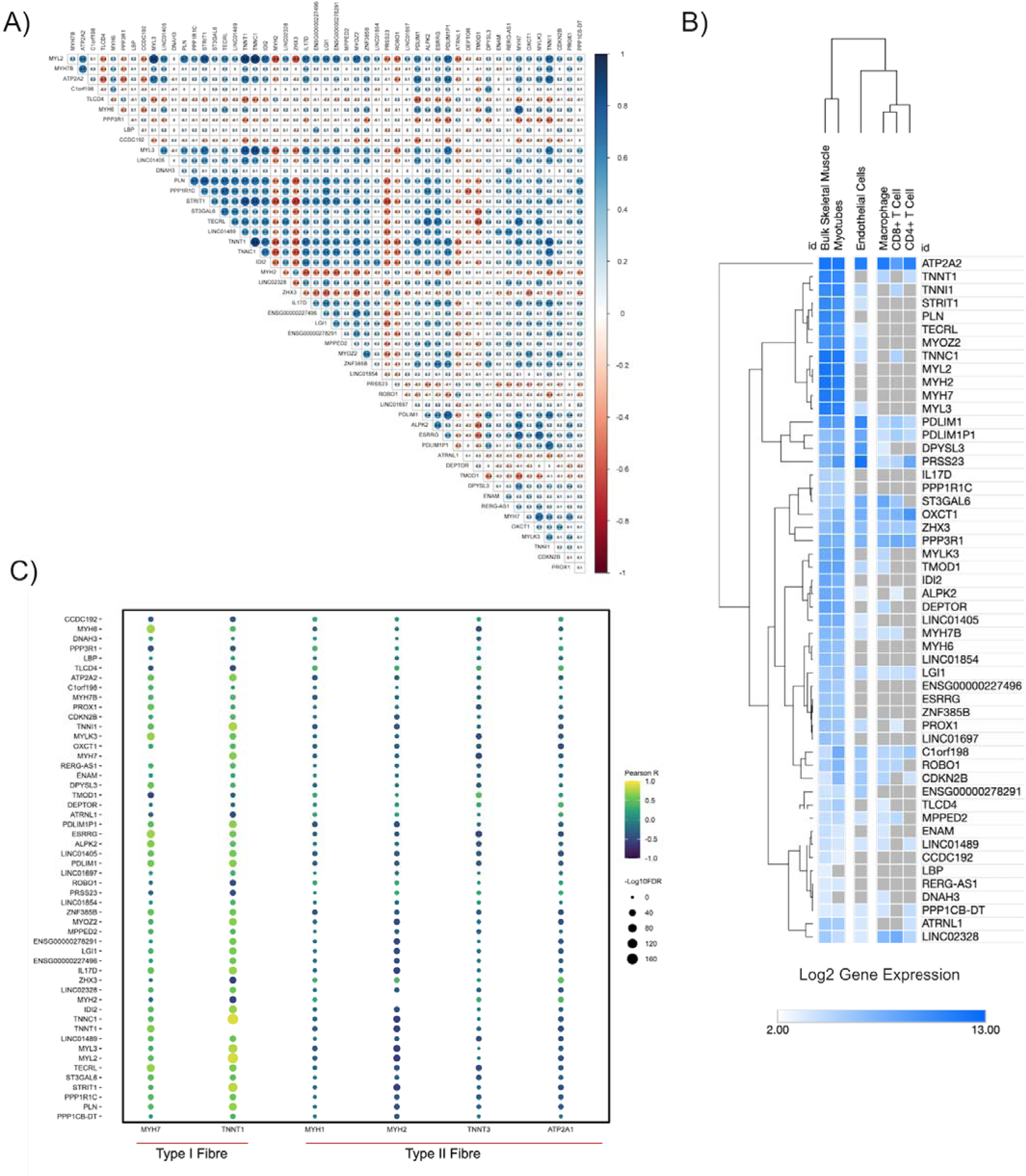
A) Pearson correlation matrix of all 52 genes co-expressed in network 2. Majority of the genes are positively correlated with each other in this network. b) the heatmap uses the log2 gene expression in skeletal muscle (n=437) and plots the 52 genes co-expressed in network 2 along with marker genes from bulk skeletal muscle, myotubes, endothelial cells, and three mononuclear cells of the immune system (macrophages, CD4 T-cells, and CD8 T-cells). The plot was created using Morpheus (https://clue.io/Morpheus), and genes and tissue types were hierarchically clustered using Euclidean distance (linkage method: complete). C) Dot plot depicting the association between the expression of network 2 genes, and gene markers for type I and type II fiber in human skeletal muscle transcriptomic data (n=437). The colouring of the dot corresponds to the Pearson correlation coefficient, and the size of the dot is proportional to the –Log^10^ FDR.

**Figure S6.**
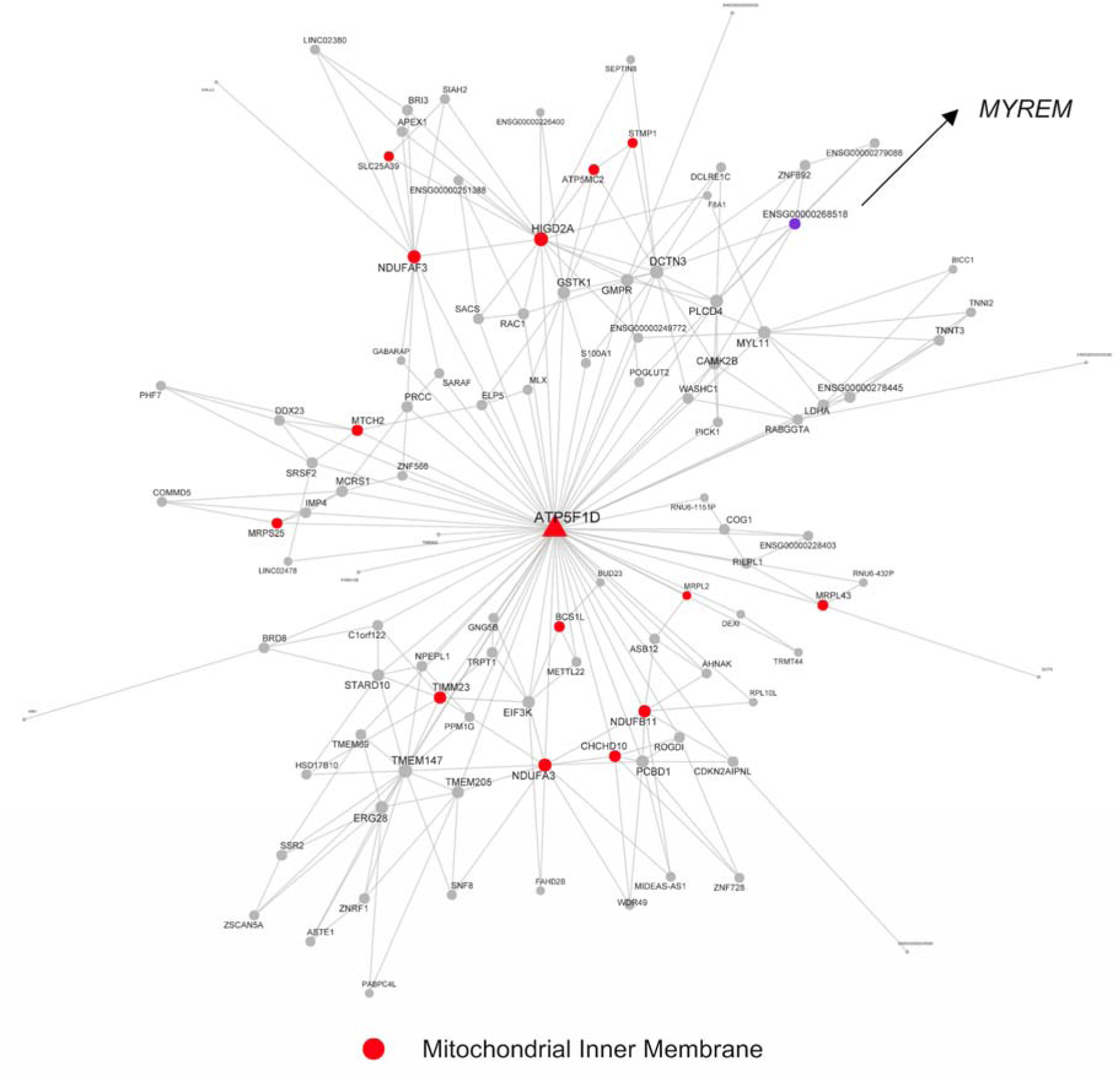
A mitochondrial-related gene co-expression network (network 4; Table 1) that contains the hypertrophy-related ncRNA gene, *MYREM* (purple).

**Figure S7.**
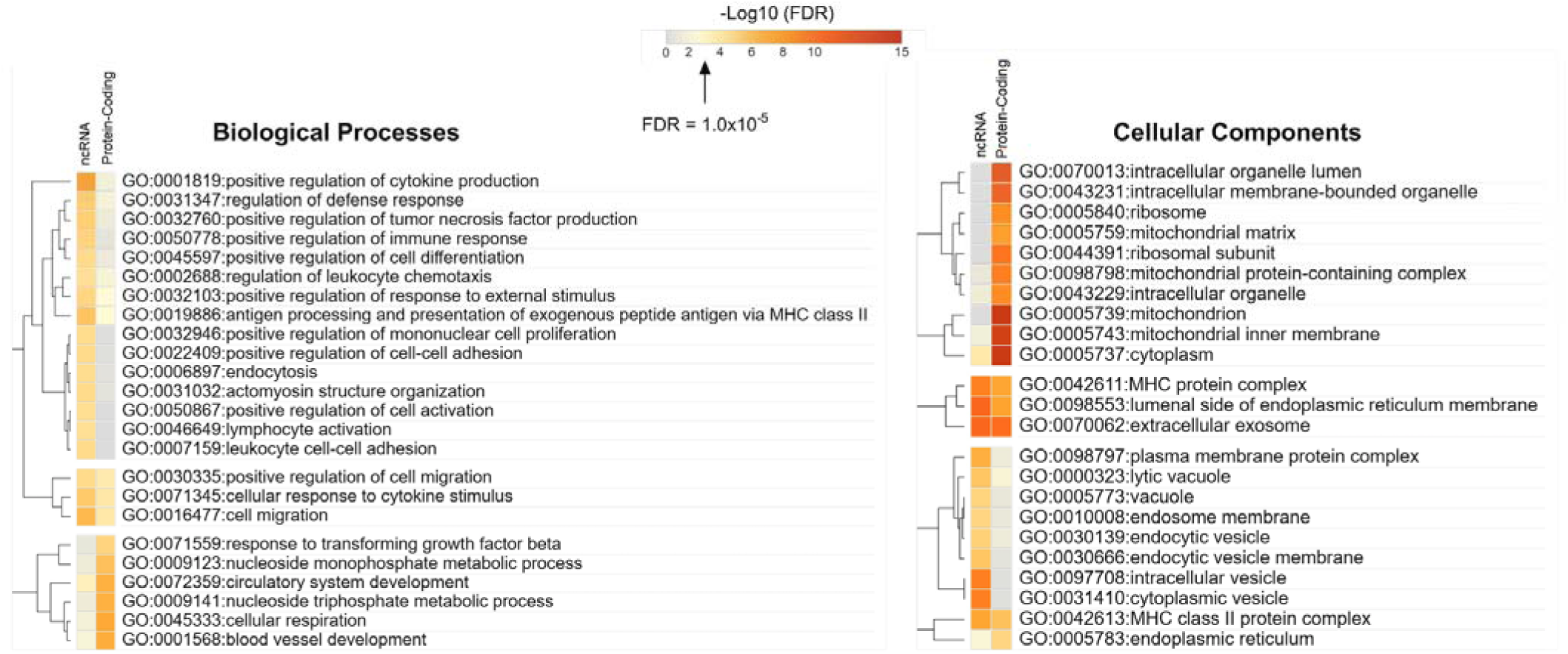
Similarities and differences in significant GO terms (biological processes and cellular components) across ncRNA MEGENA networks and our previously reported growth-regulated protein-coding MEGENA networks ^25^. The heatplot was created using Morpheus (https://clue.io/Morpheus), and genes and tissue types were hierarchically clustered using Euclidean distance (linkage method: complete).

